# Caspase Activity Affects Cell Cycle and Chromosome Integrity after Genome Editing in Human Hematopoietic Stem and Progenitor Cells

**DOI:** 10.64898/2026.11.25.769214

**Authors:** Zachary Bice, Sibabalo Sokupa, Jenree G. Maglasang, Rebekah Baskin-Doerfler, Aaron Pitre, Madison Hincher, Kalin D. Mayberry, Phillip A. Doerfler

## Abstract

Therapeutic genome editing introduces DNA double-strand breaks into hematopoietic stem and progenitor cells (HSPCs), making chromosome integrity an important consideration for edited cell function. We investigated how caspase activity influences cellular responses to genome editing in primary human HSPCs. Caspase inhibition increased the fraction of phospho-histone H3-S10-positive cells and reduced micronucleus formation and DNA damage after editing. Genome editing was associated with reduced detection of BUB1 and H2A-T120p at centromeres, with partial restoration after caspase inhibition. Genetic disruption of *BUB1* did not fully reproduce the effects of editing, and caspase inhibition continued to increase the phospho-histone H3-S10-positive fraction in *BUB1*-deficient cells. p53 suppression reduced micronucleus formation without increasing this fraction, distinguishing its effects from those of caspase inhibition. After xenotransplantation, recipients of HSPCs exposed to caspase inhibition *in vitro* had increased bone marrow cellularity and total CD34^+^ cell numbers. These findings identify caspase activity as a determinant of cell cycle and chromosome integrity outcomes after genome editing in human HSPCs.

## MAIN

Genome editing technologies precisely define where DNA double-strand breaks are introduced, but they do not control when these breaks occur within the cell cycle, creating a potential challenge for genome integrity^1,2^. The Food and Drug Administration–approved genome editing therapy for β-hemoglobinopathies (exagamglogene autotemcel) offers the prospect of lifelong symptom relief by introducing targeted DNA breaks into hematopoietic stem and progenitor cells (HSPCs) to recapitulate disease-modifying genetic variants that increase fetal hemoglobin in red blood cell progeny^3–6^. The safety and durability of a potentially lifelong therapy depend on more than repair at the intended target. Genome editing and associated *ex vivo* stress produce micronuclei and other chromosomal abnormalities, with potential consequences for the behavior and persistence of edited stem cells and the specialized cell types produced thereafter^7–12^. These outcomes depend not only on DNA repair and cell survival, but also on how cells progress through mitosis. Failure to maintain genome integrity in the stem cell pool is a defining property of bone marrow failure and cancer predisposition^13^. Understanding how edited HSPCs respond to cellular stress is therefore important for determining whether surviving cells maintain chromosomal integrity.

Cell cycle state is an important determinant of genome editing outcomes in HSPCs, particularly for precise correction through homology-directed repair (HDR). Several strategies have improved editing outcomes by controlling the timing of nuclease activity, enriching cells in HDR-permissive S/G2 phases, or coordinating transient cell cycle entry with maintenance of stem cell properties^14–19^. These studies show that editing efficiency depends not only on the targeting reagent but also on the physiological state of the cell in which editing occurs. A separate consideration is whether cells that achieve successful target modification maintain chromosome integrity during subsequent cell divisions. Cell cycle state may therefore influence genome editing outcomes through effects on chromosome segregation as well as DNA repair and stress response signaling^20,21^. Understanding these responses is necessary to evaluate the biological consequences of genome editing beyond editing efficiency alone.

Genome editing–associated chromosomal abnormalities are influenced not only by DNA repair but also by cellular responses to DNA damage and *ex vivo* culture stress. For example, extended *ex vivo* culture increases genome instability after editing^11,12^, whereas inhibition of p38 MAPK reduces micronuclei formation and improves the fitness of edited HSPCs^22,23^. The DNA damage response to genome editing in HSPCs also involves p53 activation, which can induce cell cycle arrest, apoptosis, and impair HSPC proliferation and engraftment^9,24–28^. Caspases, which mediate apoptotic responses, also have non-apoptotic functions in hematopoietic cell differentiation and maturation and have shown to regulate mitotic checkpoint proteins^29–40^. Because chromosome segregation errors can generate lagging chromosomes and micronuclei^41–43^, we asked whether caspase activity and p53 signaling influence the frequency of mitotic cells and micronucleus formation following genome editing in primary human HSPCs.

Here, we investigated how caspase activity and p53 signaling contribute to cellular responses following genome editing in primary human HSPCs. We examined cell-cycle–associated measurements, micronucleus formation, DNA damage, and centromeric BUB1 and H2A-T120p detection, and used genetic disruption of *BUB1* to assess its contribution to these responses. We further evaluated the consequences of caspase inhibition for hematopoietic recovery following xenotransplantation. These studies examine how cellular stress responses connect genome editing to chromosome integrity and the subsequent recovery of edited hematopoietic cells.

## RESULTS

### Genome editing induces persistent micronuclei in human HSPCs

DNA breaks that persist into mitosis can generate acentric chromosome fragments lacking centromeres. These fragments can missegregate during cell division and become sequestered within micronuclei^7^. Because prolonged *ex vivo* culture promotes HSPC cell cycle entry, we varied the duration of pre-stimulation before editing. To assess micronucleation after CRISPR-Cas9 editing, we used the clinically compatible MaxCyte GTx to electroporate Cas9 ribonucleoprotein (RNP) complexes in healthy-donor–derived mobilized peripheral blood CD34^+^ HSPCs with increasing times in *ex vivo* culture prior to editing of the *BCL11A* +58 enhancer (**Fig. 1a**)^3–6^. We quantified micronuclei by flow cytometry over time (**Fig. 1b–d**, **Extended Data Fig. 1a**). Genome editing increased micronuclei frequency across the experimental time course, with significantly higher micronuclei frequencies observed in edited cells for HSPCs edited immediately after thawing and after 24 or 48 hours of pre-stimulation (**Fig. 1b–d, Extended Data Fig. 1a**). The duration of pre-stimulation did not affect editing efficiency (**Fig. 1e**). Many genomic loci have been extensively validated for therapeutic editing in HSPCs. We used electroporations with RNPs to target the *HBG1/2* promoters^44,45^, *CYBB*^46^, *IL2RG*^22,23,47–50^, *CD40LG*^18,51–53^, and *WAS*^54^ in HSPCs pre-stimulated for 48 hours and observed similar patterns of editing efficiencies and rates of micronuclei (**Extended Data Fig. 1b, c**). These findings show that genome-editing–associated micronucleus formation occurs across multiple therapeutically relevant target loci and persists across a range of pre-stimulation durations despite comparable editing efficiencies.

**Figure 1.**
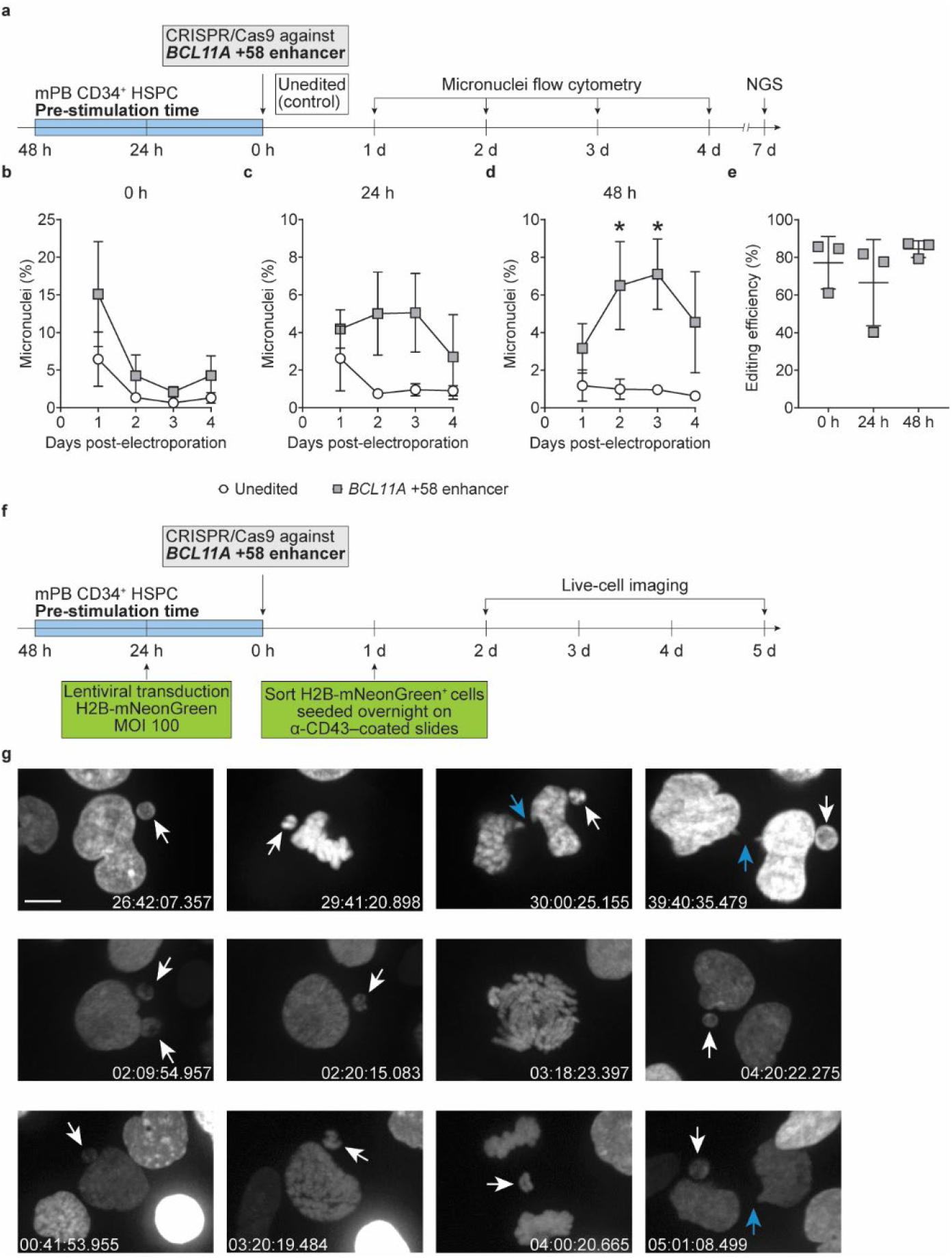
Genome editing induces micronuclei and other cytogenetic abnormalities in human HSPCs. **a,** Schematic of genome editing experiments in mobilized peripheral blood (mPB) CD34^+^ HSPCs. HSPCs were maintained for 0, 24, or 48 h before electroporation of Cas9 RNP targeting the *BCL11A* +58 enhancer using a MaxCyte GTx. **b–d,** Flow cytometry for micronuclei on days 1–4 post-electroporation in HSPCs without pre-stimulation (0 h) or with pre-stimulation for 24 or 48 h. Mean ± standard deviation with a two-way repeated measures ANOVA with the Geisser-Greenhouse correction. *n* = 3 donors performed in triplicate. Treatment effect: 0 h, *P* = 0.0030; 24 h, *P* = 0.0255; 48 h, *P* = 0.0033. Tukey’s multiple comparison test \**P* < 0.05. **e**, Genome editing efficiency of cultures in b–d determined by next-generation sequencing (NGS) on day 7 post-electroporation. Mean ± standard deviation. *n* = 3 donors performed in triplicate. **f,** Schematic of live-cell imaging experiments in mPB CD34^+^ HSPCs. **g,** Representative images of micronuclei and chromatin bridges in genome-edited HSPCs. White arrows indicate micronuclei and blue arrows indicate chromatin bridges. Scale bar = 5 μm. Timestamp shows relative time in h:m:s.ms.

The DNA contained in micronuclei can persist as micronuclei or be reincorporated into a primary nucleus during a subsequent cell division. To determine whether micronuclei-containing HPSCs persist after cell division, we performed long-term live-cell imaging of HSPCs expressing H2A-mNeonGreen to visualize chromatin dynamics, micronuclei, and mitosis (**Fig. 1f, g**; **Supplementary Videos 1–3**). Live-cell imaging showed that HSPCs containing micronuclei remained capable of progressing through subsequent cell divisions. Observed events included cells entering mitosis with visible micronuclei, changes in the number of micronuclei observed following division, and the formation of anaphase bridges. These observations show that cells containing micronuclei can remain viable and continue dividing despite persistent or additional chromosome segregation abnormalities.

There are no standard culture conditions for therapeutic genome editing of HSPCs. The experiments presented in Figure 1 were performed using culture conditions containing GlutaMAX, StemRegenin-1 (SR1), and UM729, which are commonly used to support HSPC maintenance during *ex vivo* culture^21,48,49,55–58^.

Because these additives can influence HSPC state, we next asked whether they affected cell cycle status, micronuclei formation after editing, or editing efficiency. The inclusion of GlutaMAX, SR1, and UM729 preserved quiescence, resulting in fewer cells in S and G2/M phases after 24 or 48 hours of pre-stimulation (**Extended Data Fig. 2a**). Despite the effects of these additives on cell cycle distribution, micronuclei frequencies after genome editing were comparable between the culture conditions used throughout Figure 1 and cultures lacking these additives (**Extended Data Fig. 2b–d**). Editing efficiency was similarly high without these additives (**Extended Data Fig. 2e**).

Collectively, micronuclei formation is a recurrent consequence of therapeutic genome editing in human HSPCs and micronuclei-containing cells can persist through cell division. Because p53 and apoptotic pathways shape cellular responses to DNA damage, we next tested whether their activity affects the chromosomal abnormalities that arise after genome editing. To distinguish between these possibilities, we evaluated the effects of caspase inhibition and p53 suppression on micronuclei formation and mitotic progression.

### Caspase inhibition increases the mitotic cell fraction and reduces micronucleus formation in cycling HSPCs

Because pre-stimulation changes the cell cycle state of HSPCs, we evaluated the effects of caspase inhibition on micronucleus formation in cells edited immediately after thawing or following 48 hours of pre-stimulation. In quiescent HSPCs, micronuclei frequencies were highest 1 day after electroporation and declined thereafter, with little effect of Q-VD-OPh treatment (**Fig. 2a**). In contrast, after 48 hours of pre-stimulation, Q-VD-OPh treatment reduced editing-associated micronuclei formation toward levels observed in unedited cells (**Fig. 2b**).

**Figure 2.**
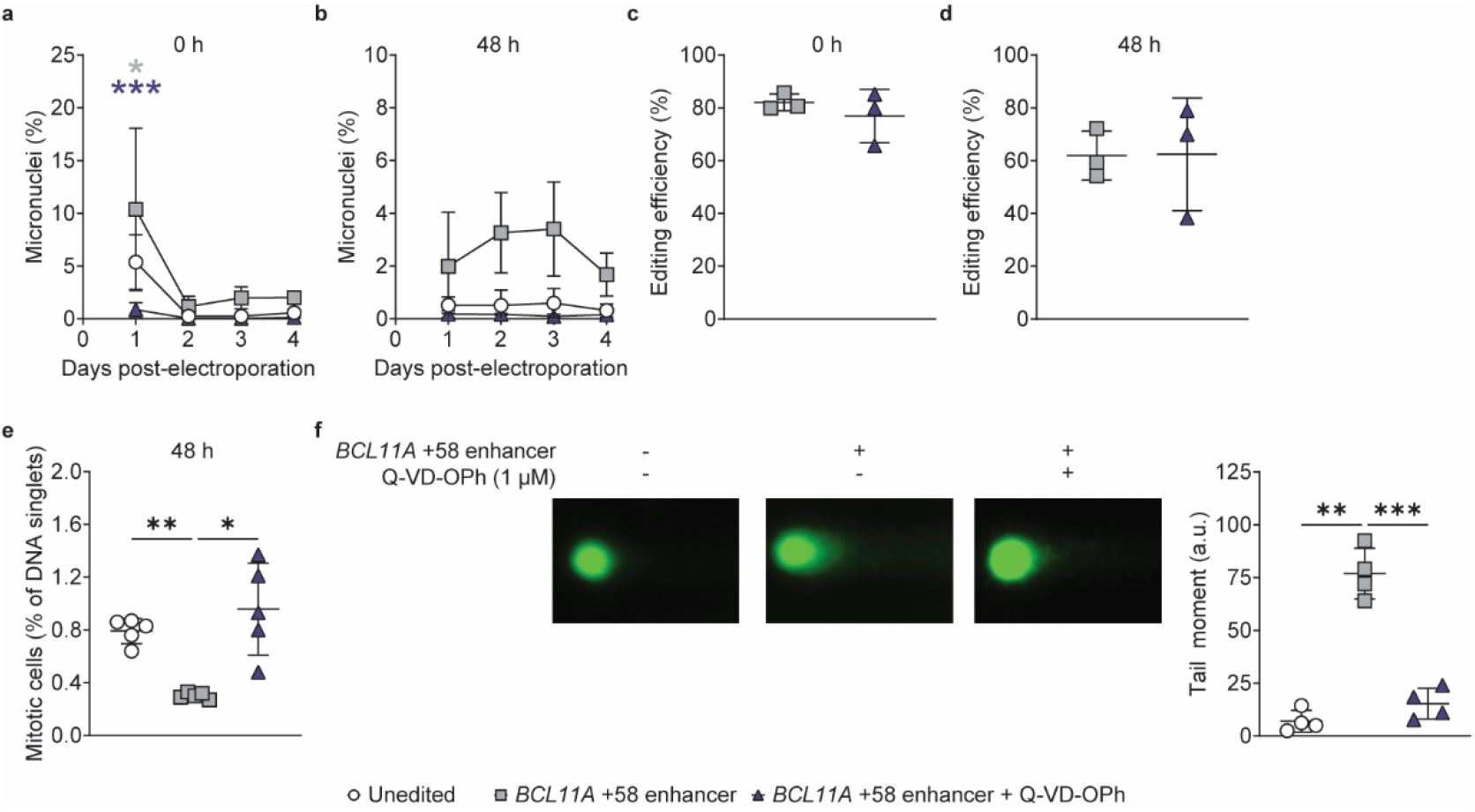
Caspase inhibition alters micronucleus formation, mitotic cell representation, and DNA damage after genome editing. Flow cytometry for micronuclei on days 1–4 post-electroporation in HSPCs without pre-stimulation (0 h) ± caspase inhibition (Q-VD-OPh). Mean ± standard deviation with a two-way repeated measures ANOVA with the Geisser-Greenhouse correction. *n* = 3 donors performed in triplicate. Time effect: *P* = 0.0003. Treatment effect: *P* = 0.052. Tukey’s multiple comparison test \**P* < 0.05, \*\*\**P* < 0.001. Grey * indicates significant differences between unedited and edited. Blue *** indicates significant differences between edited and edited + Q-VD-OPh. **b,** Flow cytometry for micronuclei on days 1–4 post-electroporation in HSPCs with 48 h pre-stimulation ± caspase inhibition. Mean ± standard deviation with a two-way repeated measures ANOVA with the Geisser-Greenhouse correction. *n* = 3 donors performed in triplicate. Treatment effect: *P* = 0.0033. **c, d,** Genome editing efficiency of cultures in a, b determined by next-generation sequencing on day 7 post-electroporation. Mean ± standard deviation. *n* = 3 donors performed in triplicate. **e,** Flow cytometry analysis of the fraction of phospho-histone H3-S10-positive cells 1 day post-electroporation in HSPCs with 48 h pre-stimulation. Mean ± standard deviation with a one-way repeated measures ANOVA with the Geisser-Greenhouse correction. *n* = 5 donors. Holm-Šídák’s multiple comparisons test \**P* < 0.05, \*\**P* < 0.01. **f,** Tail moment measurements from alkaline comet assay 1 day post-electroporation in HSPCs with 48 h pre-stimulation. Mean ± standard deviation with a one-way repeated measures ANOVA with the Geisser-Greenhouse correction. *n* = 5 donors. Each dot represents the donor-level average tail moment. The total number of nuclei analyzed was 226 for unedited, 258 for edited, and 250 for edited + Q-VD-OPh. Holm-Šídák’s multiple comparisons test \*\**P* < 0.01, \*\*\**P* < 0.001.

Caspase inhibition reduced editing-associated micronuclei formation without affecting editing efficiency (**Fig. 2c, d**). As expected, Q-VD-OPh reduced apoptotic cell death after editing (**Extended Data Fig. 3a, b**). Because caspase inhibition reduced both micronuclei formation and apoptotic cell death after genome editing, we next asked whether these effects were associated with alterations in cell cycle progression. Genome editing produced smaller changes in cell cycle distribution under quiescent conditions than after 48 h pre-stimulation (**Extended Data Fig. 3c**). To distinguish effects of editing from those of pre-stimulation, cell cycle distributions were compared between edited and unedited cells cultured under identical conditions. After 48 hours of pre-stimulation, edited cells consistently showed lower G1 fractions and higher S and G2/M fractions than matched unedited controls throughout the observation period (**Extended Data Fig. 3d**). Because DNA content analysis cannot distinguish cells arrested in G2 from those progressing through mitosis, we measured phospho-histone H3-S10, a marker of mitotic chromatin. Genome editing reduced the fraction of phospho-histone H3-S10-positive cells, whereas caspase inhibition increased this fraction (**Fig. 2e**). These measurements indicate that caspase inhibition increases the representation of mitotic cells at the time of analysis but do not establish the rate of mitotic entry or the fidelity of chromosome segregation. Q-VD-OPh also reduced DNA damage measured by alkaline comet assay (**Fig. 2f**). These data suggest the increase in the fraction of phospho-histone H3-S10-positive cells occurred alongside a reduction in detectable DNA damage, rather than an increase in DNA damage.

### p53 suppression reduces micronucleus formation without increasing the mitotic cell fraction

Given the activation of p53-dependent DNA damage responses after genome editing^9,24,25,48,59^, we examined whether suppression of p53 alters genome-editing–associated micronucleus formation and mitotic progression. HSPCs were edited at the *BCL11A* +58 enhancer alone or simultaneously with *TP53* or co-electroporated with the dominant-negative p53 inhibitor GSE56 (**Extended Data Fig. 4a, b**)^47,48,60^. In HSPCs edited without pre-stimulation, micronuclei frequencies declined over time irrespective of treatment (**Extended Data Fig. 4c**). After 48 hours of pre-stimulation, genome editing increased micronuclei frequency, and both *TP53* disruption and GSE56 expression were associated with reduced micronuclei frequencies relative to edited controls (**Extended Data Fig. 4d**). Despite effects on micronuclei formation, neither *TP53* disruption nor GSE56 expression increased the proportion of phospho-histone H3-S10-positive cells after genome editing (**Extended Data Fig. 4e**). These findings show that p53 suppression can reduce micronucleus formation without increasing the fraction of phospho-histone H3-S10-positive cells, distinguishing its effects from those of caspase inhibition.

### Genome editing is associated with reduced detection of BUB1 at centromeres

We next investigated whether caspase activity influences mitotic regulators involved in chromosome segregation. BUB1 is a key regulator of chromosome segregation and mitotic checkpoint signaling and has been reported to undergo caspase-dependent cleavage^38–40^. To confirm this relationship, we treated HL-60 cells with etoposide, which induced CASPASE-3 activation and generation of BUB1 cleavage products (**Extended Data Fig. 5a, b**). Genetic disruption of *CASP3* prevented BUB1 cleavage after etoposide treatment, confirming BUB1 as a CASPASE-3 substrate in this system (**Extended Data Fig. 5c, d**).

We next examined whether genome editing altered BUB1-associated signaling in primary human HSPCs. To assess centromeric BUB1 signaling, we evaluated both BUB1 localization and phosphorylation of histone H2A at threonine 120 (H2A-T120p), a target of BUB1 kinase activity at centromeres^61^. Genome editing reduced pro-CASPASE-3 abundance, consistent with caspase activation, and this effect was reversed by Q-VD-OPh treatment (**Fig. 3a, b**). Although BUB1 cleavage was readily detected after etoposide treatment in HL-60 cells, total BUB1 protein abundance in genome-edited HSPCs varied among donors and did not differ significantly between conditions (**Fig. 3c, d**). Analysis of centromeric signaling identified lower frequencies of BUB1-positive, H2A-T120p-positive, and BUB1/H2A-T120p double-positive centromeres in several donors after editing, with partial restoration after caspase inhibition (**Fig. 3e–k**). Fluorescence intensity among positive centromeres was comparatively preserved. Together, these findings indicate that genome editing was associated with a reduction in the frequency of centromeres with detectable BUB1 and H2A-T120p staining, rather than a uniform reduction in fluorescence intensity across positive centromeres. Because the BUB1 antibody recognizes both full-length and cleaved BUB1, these measurements do not distinguish changes in BUB1 abundance from changes in its proteolytic processing.

**Figure 3.**
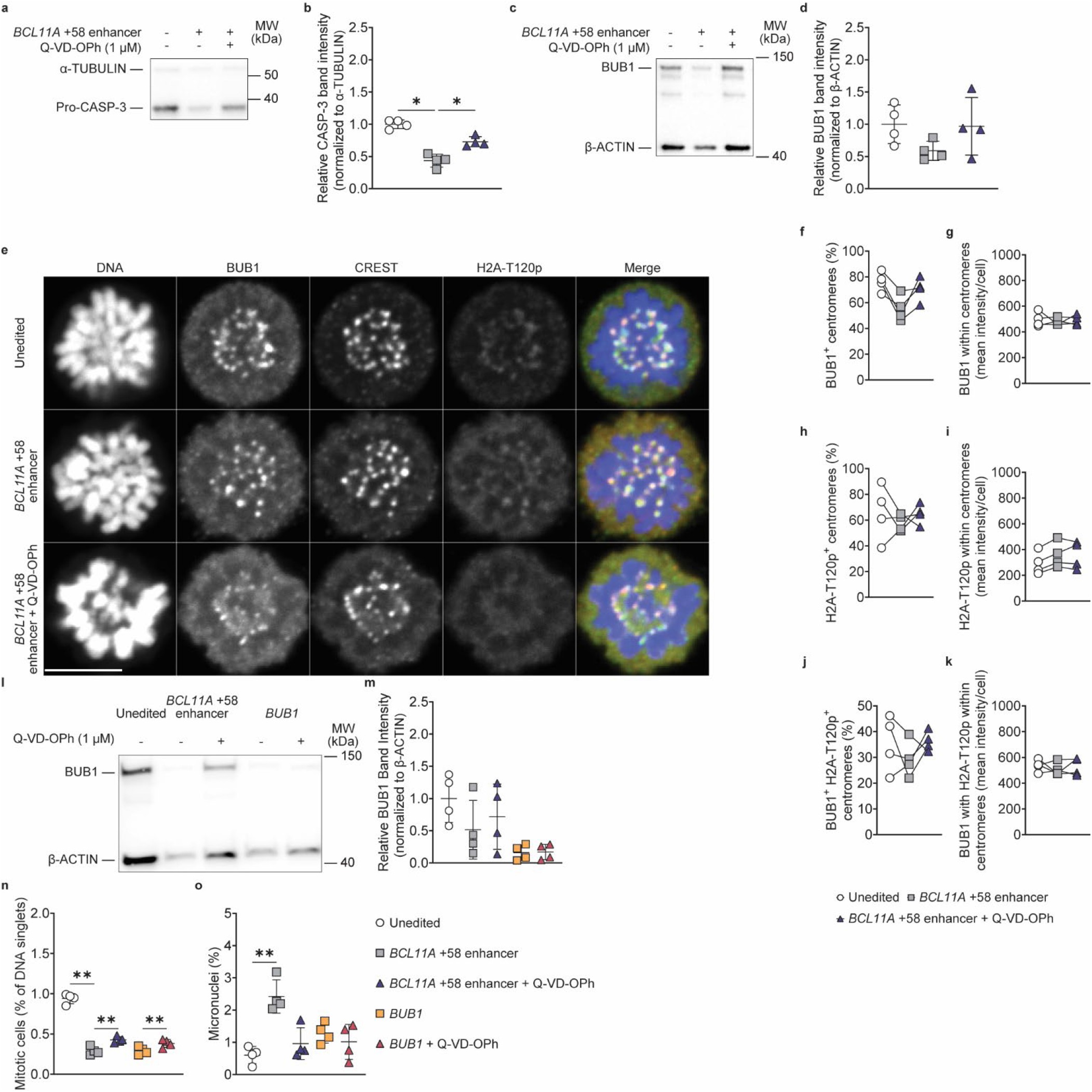
Genome editing alters centromeric BUB1 and H2A-T120p detection. **a,** Representative immunoblot for CASPASE-3 two days post-electroporation in HSPCs with 48 h pre-stimulation. **b,** Quantification of pro-CASPASE-3 normalized to α-TUBULIN. Mean ± standard deviation with a one-way repeated measures ANOVA with the Geisser-Greenhouse correction. *n* = 4 donors. Holm-Šídák’s multiple comparisons test \**P* < 0.05. **c,** Representative immunoblot for BUB1 two days post-electroporation in HSPCs with 48 h pre-stimulation. **d,** Quantification of BUB1 normalized to β-ACTIN. Mean ± standard deviation. *n* = 4 donors. **e,** Representative confocal images of unedited HSPCs or 1 day post-electroporation with Cas9 RNP targeting the *BCL11A* +58 enhancer ± Q-VD-OPh. In the merged panel, DNA (blue), BUB1 (green), centromere (CREST, orange), and histone H2A phosphorylated at threonine 120 (H2A-T120p, red) are shown. Scale bar = 5 μm. **f,** Quantification of BUB1^+^ centromere frequency. **g,** Quantification of BUB1 mean fluorescence intensity at centromeres per cell. **h,** Quantification of H2A-T120p^+^ centromere frequency. **i,** Quantification of H2A-T120p mean fluorescence intensity at centromeres per cell. **j,** Quantification of BUB1^+^H2A-T120p^+^ double-positive centromere frequency. **k,** Quantification of BUB1 H2A-T120pdouble-positive mean fluorescence intensity at centromeres per cell. *n* = 4 donors. For f–k, each point represents an individual donor with lines connecting paired conditions. **l,** Immunoblot for BUB1 and β-ACTIN in unedited HSPCs or 2 days post-electroporation with Cas9 RNP targeting the *BCL11A* +58 enhancer or *BUB1* ± Q-VD-OPh. **m,** Quantification of BUB1 normalized to β-ACTIN. Mean ± standard deviation. *n* = 4 donors. **n,** Flow cytometry analysis for mitotic cells (phospho-H3-S10) in unedited HSPCs or 1 day post-electroporation with Cas9 RNP targeting the *BCL11A* +58 enhancer or *BUB1* ± Q-VD-OPh. Mean ± standard deviation with a one-way repeated measures ANOVA with the Geisser-Greenhouse correction. *n* = 5 donors. Holm-Šídák’s multiple comparisons test \*\**P* < 0.01. **o,** Flow cytometry for micronuclei in unedited HSPCs or 2 days post-electroporation with Cas9 RNP targeting the *BCL11A* +58 enhancer or *BUB1* ± Q-VD-OPh. Mean ± standard deviation with a one-way repeated measures ANOVA with the Geisser-Greenhouse correction. *n* = 4 donors. Holm-Šídák’s multiple comparisons test \*\**P* < 0.01.

To determine whether BUB1 contributes functionally to genome-editing–associated mitotic phenotypes, we directly disrupted *BUB1* in HSPCs. Editing of *BUB1* resulted in near-complete loss of BUB1 protein regardless of Q-VD-OPh treatment, confirming efficient target depletion (**Fig. 3l, m**). Similar to *BCL11A* +58 enhancer editing, *BUB1* disruption reduced the frequency of phospho-histone H3-S10-positive cells and increased micronuclei formation (**Fig. 3n, o**). However, these effects were less pronounced than those observed after *BCL11A* +58 enhancer editing. Furthermore, Q-VD-OPh continued to increase the fraction of phospho-histone H3-S10-positive cells in *BUB1*-disrupted cultures. Thus, although *BUB1* disruption reduced the fraction of mitotic cells, loss of BUB1 did not reproduce the full phenotype associated with genome editing, and the effect of caspase inhibition on the mitotic cell fraction did not require detectable BUB1 protein.

### Caspase inhibition increases early hematopoietic cell recovery after transplantation

Having established that caspase inhibition reduced micronuclei formation and increased phospho-histone H3-S10-positive HSPCs after genome editing, we next investigated whether transient caspase inhibition influences the *in vivo* function of edited HSPCs. To address this question, we cultured HSPCs edited at the *BCL11A* +58 enhancer with Q-VD-OPh for 24 hours immediately after electroporation until xenotransplantation into NBSGW mice (**Fig. 4a**)^54^. The transplantation study was designed to assess early hematopoietic output rather than long-term stem cell self-renewal or competitive repopulation activity.

**Figure 4.**
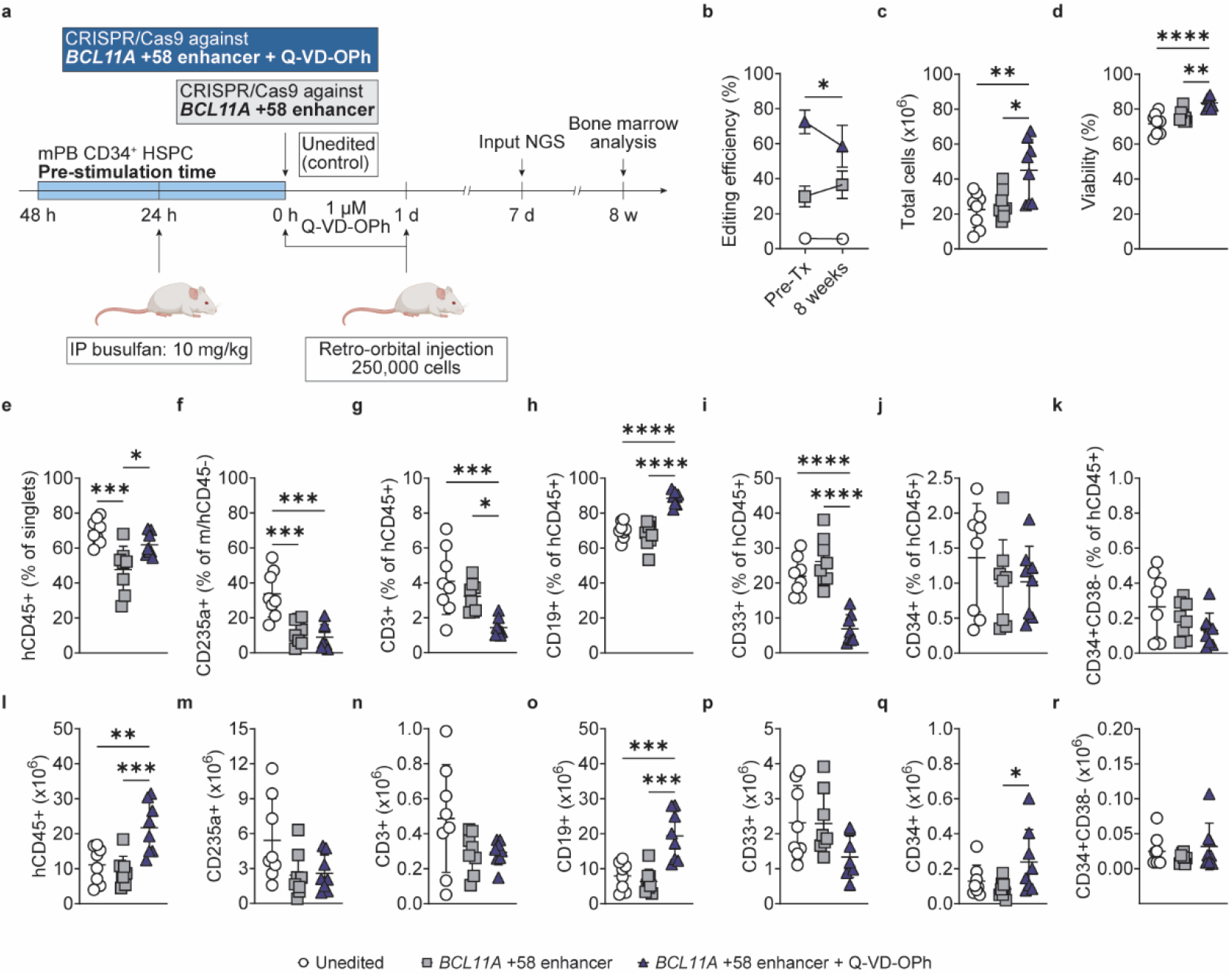
Caspase inhibition increases hematopoietic cell recovery after xenotransplantation. **a,** Schematic of genome editing and xenotransplantation of CD34^+^ HSPCs into immunodeficient NBSGW mice. HSPCs electroporated with Cas9 RNP targeting the *BCL11A* +58 enhancer were cultured in 1 μM Q-VD-OPh for 24 h immediately after electroporation until injection into NBSGW mice. IP, intraperitoneal. **b,** Genome editing efficiency of the input HSPCs collected on day 7 post-electroporation and whole bone marrow cells after 8 weeks determined by next-generation sequencing (NGS). Mean ± standard deviation with a two-tailed, unpaired t-test. *n* = 5 (four individual donors and the pooled cells for injection) for pre-transplant (Pre-Tx) and *n* = 8 mice/group at 8 weeks, \**P* < 0.05 between pre-transplant and 8 weeks post-transplant in the Q-VD-OPh treated groups. **c,** Total number of nucleated cells collected from recipient mouse bone marrow. **d,** Viability of nucleated cells collected from recipient mouse bone marrow. **e,** Frequency of human (h) CD45^+^ within the singlet population from recipient mouse bone marrow. **f,** Frequency of CD235a^+^ erythroid cells within the mouse (m) and hCD45 double-negative population. **g,** Frequency of CD3^+^ T cells within the hCD45^+^ population. **h,** Frequency of CD19^+^ B cells within the hCD45^+^ population. **i,** Frequency of CD33^+^ myeloid cells within the hCD45^+^ population. **j,** Frequency of CD34^+^ cells within the hCD45^+^ population. **k,** Frequency of CD34^+^CD38^-^ cells within the hCD45^+^ population. **l,** The total number of hCD45^+^ from recipient mouse bone marrow. **m,** The total number of CD235a^+^ erythroid cells from recipient mouse bone marrow. **n,** The total number of CD3^+^ T cells from recipient mouse bone marrow. **o,** The total number of CD19^+^ B cells from recipient mouse bone marrow. **p,** The total number of CD33^+^ myeloid cells from recipient mouse bone marrow. **q,** The total number of CD34^+^ cells from recipient mouse bone marrow. **r,** The total number of CD34^+^CD38^-^ cells from recipient mouse bone marrow. For c–r, mean ± standard deviation with an ordinary one-way ANOVA. *n* = 8 mice/group. Tukey’s multiple comparisons test \**P* < 0.05, \*\**P* < 0.01, \*\*\**P* < 0.001, \*\*\*\**P* < 0.0001.

Edited alleles remained detectable in human cells recovered from all recipient mice after transplantation, although editing frequencies in the recipients of Q-VD-OPh HSPCs were lower than those measured in the pre-transplant samples (**Fig. 4b**). Recipient mice transplanted with Q-VD-OPh-treated HSPCs had higher bone marrow cellularity and higher numbers of viable recovered cells compared with mice receiving unedited or edited HSPCs without caspase inhibition (**Fig. 4c, d**). Human hematopoietic cell engraftment was also improved with Q-VD-OPh treatment in comparison to editing alone, as indicated by a higher frequency of hCD45^+^ cells within recipient bone marrow (**Fig. 4e**). Analysis of lineage output showed multilineage hematopoiesis in all groups, including erythroid (CD235a^+^), T cell (CD3^+^), B cell (CD19^+^), myeloid (CD33^+^), and CD34^+^ populations (**Fig. 4f–k**). Consistent with the higher cellularity of the recipient bone marrow, Q-VD-OPh-treated grafts yielded greater absolute recovery of human hematopoietic cells (**Fig. 4l–r**). Significantly more cells were recovered within the hCD45^+^, CD19^+,^ and CD34^+^ compartments (**Fig. 4l, o, r**). While multilineage hematopoiesis was maintained in all groups, Q-VD-OPh treatment was associated with changes in lineage distribution, indicating that increased cellular output was not evenly distributed across hematopoietic lineages. These findings indicate that transient caspase inhibition enhances hematopoietic recovery from genome edited HSPCs without compromising editing efficiency.

## DISCUSSION

Our findings show that caspase activity influences several distinct cellular outcomes following genome editing in primary human HSPCs, including the fraction of cells positive for phospho-histone H3-S10, micronucleus formation, and hematopoietic cell recovery after transplantation (**Fig. 5**). Genome editing can produce a range of genomic alterations beyond those at the intended target, including large-scale chromosomal rearrangements and copy-number changes, and these outcomes have been linked to DNA repair and cellular responses to editing-induced damage^1,2,7–12^. Our results add to this work by showing that caspase inhibition changes both the representation of mitotic cells and the frequency of micronuclei after editing. These findings emphasize that editing efficiency and cell survival alone do not capture all the cellular consequences of genome editing.

**Figure 5.**
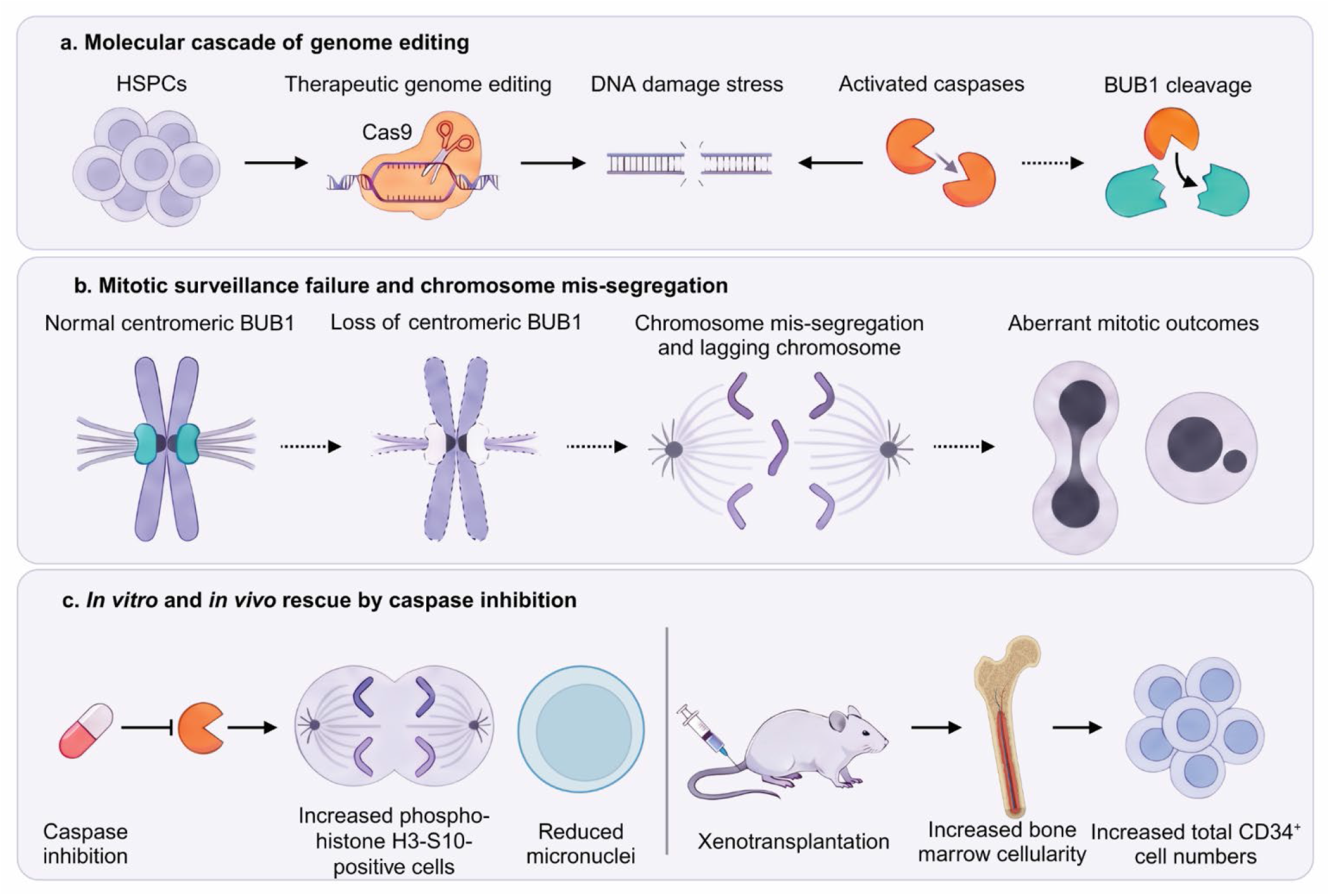
Model of caspase-associated cellular outcomes after genome editing. **a,** Genome editing induces cellular stress and caspase activation in hematopoietic stem and progenitor cells (HSPCs). Caspase-associated BUB1 cleavage is a potential link between cellular stress and altered mitotic regulation. **b,** Genome editing is associated with reduced frequency of centromeres with detectable BUB1 and H2A-T120p. These changes may contribute to altered mitotic surveillance and chromosome segregation, although a direct causal relationship has not been established. **c,** Caspase inhibition increases the fraction of phospho-histone H3-S10-positive cells and reduces micronucleus formation *in vitro*. The transient inhibition of caspases before xenotransplantation resulted in increased bone marrow cellularity and total CD34^+^ cell numbers after 8 weeks. These findings support the evaluation of mitotic integrity alongside cell viability and hematopoietic output in the assessment of the preclinical safety of genome-editing approaches. Solid arrows indicate directly studied pathways, whereas dashed arrows indicate proposed mechanisms that have not been directly established in primary HSPCs. Created in BioRender. https://BioRender.com/w26vhkn.

A notable finding was that caspase inhibition increased the fraction of phospho-histone H3-S10-positive cells while reducing micronucleus formation. This result argues against the possibility that the lower frequency of micronuclei simply reflects a reduced representation of mitotic cells after treatment. However, phospho-histone H3-S10 measurements provide a snapshot of cells in mitosis and do not establish the rate of mitotic entry, the duration of mitosis, or the fidelity of chromosome segregation. Similarly, micronucleus measurements do not identify the specific division in which a micronucleus formed. Because chromosome segregation errors can generate lagging chromosomes and micronuclei, these findings raise the possibility that caspase activity influences processes associated with chromosome integrity during cell division^38–40^. The reduction in DNA damage detected by alkaline comet assay after Q-VD-OPh treatment also suggests that changes in DNA damage may contribute to the reduction in micronucleus formation. Thus, the effects of caspase inhibition on micronuclei cannot be attributed specifically to altered chromosome segregation.

The effects of p53 suppression and caspase inhibition provide insight into distinct aspects of the cellular response to genome editing. Both *TP53* disruption and GSE56 expression reduced micronucleus formation, while caspase inhibition also increased the fraction of phospho-histone H3-S10-positive cells. This distinction is relevant because p53 signaling is an established component of the cellular response to CRISPR-Cas9 editing and can influence cell cycle arrest, DNA repair, and the propagation of cells with chromosomal abnormalities^9,24,26–28^. The stimulus responsible for caspase activation in edited HSPCs remains to be determined and may involve electroporation and RNP delivery, Cas9-induced DNA breaks, or a combination of these stresses. Our findings place caspase activity alongside p53 signaling as a contributor to cellular outcomes after editing, with each pathway associated with a distinct pattern of effects across the measured endpoints.

Genome editing was also associated with changes in centromeric BUB1 and H2A-T120p detection. BUB1 participates in spindle assembly checkpoint signaling and chromosome congression, and can undergo caspase-dependent cleavage during apoptosis^39,40^. BUB1-dependent phosphorylation of H2A-T120 contributes to centromeric recruitment of shugoshin and chromosome stability^61^. In our study, editing reduced the frequency of centromeres with detectable BUB1 and H2A-T120p staining, with partial restoration after caspase inhibition. These observations connect caspase activity with changes in the detection of centromeric proteins involved in chromosome regulation during mitosis. Total BUB1 protein abundance did not differ significantly between conditions, and our immunofluorescence measurements could not distinguish full-length BUB1 from cleavage products. Genetic disruption of *BUB1* also did not fully reproduce the effects of genome editing, and Q-VD-OPh continued to increase the fraction of phospho-histone H3-S10-positive cells in BUB1-disrupted cultures. Thus, changes in centromeric BUB1 detection may contribute to the cellular response to editing, while the effects of caspase inhibition on the mitotic cell fraction likely involve additional targets or processes.

Caspases also have functions beyond their roles in apoptosis, including contributions to hematopoietic differentiation and maturation^29–37^. These activities provide a broader context for interpreting the effects of caspase inhibition in edited HSPCs, particularly because Q-VD-OPh altered both cell cycle-associated measurements and hematopoietic output after xenotransplantation. Caspase-dependent cleavage of mitotic regulators has also been reported in other systems^33,34,36,39^, providing a potential link between caspase activity and the cellular changes observed here. Our findings do not identify the relevant caspase substrates in primary HSPCs or establish direct cleavage of BUB1 in this setting. The continued effect of Q-VD-OPh on the phospho-histone H3-S10-positive fraction in *BUB1*-disrupted cells is consistent with caspase-dependent effects extending beyond BUB1.

The duration of *ex vivo* pre-stimulation also influenced the effects of caspase inhibition on micronucleus formation. Q-VD-OPh had little effect on micronucleus frequency in cells edited immediately after thawing, whereas it reduced micronucleus formation in cells edited after 48 hours of pre-stimulation. *Ex vivo* culture can alter HSPC activity and influence the outcomes of genome editing and transplantation^11,12,20,21^. In this context, our findings suggest that the effects of caspase inhibition depend on the cellular state established during *ex vivo* culture. Because pre-stimulation duration and cell cycle state were not independently varied in our experiments, the contribution of cell cycle state to this difference remains to be determined.

Finally, transient caspase inhibition increased bone marrow cellularity and the absolute numbers of recovered human hematopoietic cells, including CD34^+^ cells, 8 weeks after transplantation. These findings indicate increased early hematopoietic cell recovery under the conditions tested, but they should not be interpreted as evidence that genomic abnormalities are eliminated or that stem cell function is improved. The lower editing frequencies in recovered cells relative to pre-transplant samples also indicate that the representation of edited alleles changed during xenotransplantation. The basis for this change was not determined. More broadly, these results emphasize that cellular recovery, editing frequency, and chromosome integrity are distinct outcomes that should be evaluated separately when assessing the consequences of genome editing.

Together, these findings show that caspase activity influences cell cycle-associated measurements, micronucleus formation, and early hematopoietic recovery following genome editing in primary human HSPCs. The effects on these outcomes are not fully explained by p53 signaling or BUB1, and the relationship between the observed changes in mitotic cell fraction, DNA damage, and micronucleus formation remains to be resolved. Evaluating these outcomes separately will be important for understanding the cellular consequences of therapeutic genome editing.

## LIMITATIONS OF THE STUDY

First, the experiments do not establish the stimulus responsible for caspase activation after genome editing. Because the study did not distinguish the effects of electroporation and RNP delivery from those of Cas9-induced DNA double-strand breaks, we cannot determine whether caspase activation is a result of the editing procedure itself, by DNA damage, or by a combination of these stresses. The distinct effects of p53 suppression and caspase inhibition on the fraction of phospho-histone H3-S10-positive cells indicate that these pathways have different effects on the measured cellular outcomes, but do not establish the stimulus responsible for caspase activation.

Second, our measurements do not establish the mechanism by which caspase inhibition reduces micronucleus formation. Phospho-histone H3-S10 staining provides a population-level measurement of cells in mitosis at a defined time, whereas micronucleus frequency was measured at a separate time point. Because micronuclei occurred in a minority of cells within a heterogeneous primary HSPC population, these measurements do not establish whether cells with altered cell cycle behavior are the same cells that subsequently develop micronuclei. In addition, caspase inhibition reduced DNA damage measured by alkaline comet assay, and this reduction may contribute to the lower micronucleus frequency. The experiments also do not distinguish altered chromosome segregation from changes in the survival or persistence of cells containing chromosomal abnormalities.

Third, although centromeric BUB1 and H2A-T120p measurements were consistent with reduced detection of these proteins after genome editing, not all measures reached statistical significance and substantial donor-to-donor variability was observed. Total BUB1 protein abundance did not differ significantly between conditions, and our immunofluorescence measurements could not distinguish full-length BUB1 from potential cleavage products. We did not directly assess BUB1 proteolysis in genome edited primary HSPCs. Furthermore, genetic disruption of *BUB1* did not fully reproduce the editing-associated phenotypes, and caspase inhibition continued to increase the fraction of H3-S10-positive cells in BUB1-deficient cultures. The contribution of BUB1 processing to these observations, along with the identity of additional relevant caspase substrates, remains to be determined.

Fourth, the effects of caspase inhibition on micronucleus formation depended on pre-stimulation history. Although the pre-stimulation conditions differed in cell cycle distribution, they also differed in the duration of *ex vivo* culture. We therefore cannot distinguish the contribution of cell cycle state from other culture-associated changes in HSPC physiology. The study did not directly assess cellular senescence or other features of HSPC state that could influence the response to editing.

Finally, the effects of caspase inhibition on post-transplantation bone marrow cellularity and total human CD34^+^ cell numbers do not establish improved long-term stem cell function or genomic integrity. Because recipients were analyzed 8 weeks after transplantation of relatively low numbers of edited mobilized peripheral blood CD34^+^ cells, the observed increases in human cell recovery are most consistent with effects on early hematopoietic output rather than durable stem cell function. Long-term and secondary transplantation studies will be required to determine whether these changes persist and affect durable HSC function. The present study also examined a specific genome-editing platform and primary HSPC culture system. The extent to which similar effects occur across other editing methods, disease settings, or hematopoietic populations remains to be determined.

## METHODS

All research complies with the St. Jude Children s Research Hospital (St. Jude) and Medical College of Wisconsin (MCW) Institutional Biosafety Committees, the MCW Institutional Animal Care and Use Committee, and was performed according to relevant ethical regulations. The use of de-identified primary cells described below was considered non-human subjects research and exempt by St. Jude and MCW Institutional Review Boards.

### Cell culture

Human promyelocytic acute myeloid leukemia cells were purchased from ATCC (CCL-240, HL-60). HL-60 cells were maintained in IMDM (Thermo, 12440053) supplemented with 10% FBS (Thermo, 26140-079), 100 U/mL penicillin, and 100 μg/mL streptomycin (Thermo, 15140163). Etoposide was purchased from Selleckchem (S1225).

Human CD34^+^ HSPCs were purchased from StemExpress or the Hematopoietic Cell Procurement and Processing Resource at Fred Hutchinson Cancer Center. After informed consent was acquired, CD34^+^ HSPCs were isolated from healthy male and female donor peripheral blood after mobilization and enriched by immunomagnetic bead selection using an AutoMACS instrument (Miltenyi Biotec). Cryopreserved HSPCs were thawed and pre-stimulated for 0, 24, or 48 hours in X-VIVO 10 (Lonza, BP04-743Q) with 100 ng/mL SCF (R&D Systems, 255-SC), FLT3-L (R&D Systems, 308-FK), and TPO (R&D Systems, 288-TPH).

Where indicated, HSPCs were supplemented with 1x GlutaMAX (Thermo, 35050-061), 0.5 nM UM729 (StemCell Technologies, 72332), 1 μM StemRegenin (SR)-1 (StemCell Technologies, 72344), and 1 μM Q-VD-OPh (Selleckchem, S7311). All cell culture was performed at 37 °C with 5% CO_2_ in a water-jacketed incubator.

### Genome editing and editing efficiency

HL-60 cells were electroporated using a Neon electroporation system (Thermo). HSPCs were electroporated using Neon or MaxCyte GTx. Purified recombinant Cas9 protein (3x-NLS) was obtained from the St. Jude Protein Production Facility. Chemically modified single guide RNAs (sgRNA) were synthesized by Synthego with 2′-O-methyl 3′-phosphorothioate modifications between the three terminal nucleotides at both the 5′ and 3′ ends. RNP complexes were formed by incubating Cas9 (5 μM) with sgRNAs (15 μM) in manufacturer–approved buffers for at least 30 minutes. Cells were washed in DPBS, resuspended in the manufacturer–provided buffers for cell lines or primary cells (Thermo Fisher Scientific), mixed with RNP complexes, and electroporated using program 5 (HL-60) or program 24 (HSPC). For MaxCyte electroporations, RNP molar ratios were maintained, HSPCs were washed in DPBS, resuspended in the manufacturer–approved buffer (MaxCyte, EPB1), mixed with RNP complexes, and electroporated using program HSC-3 after 0, 24, or 48 hours of pre-stimulation. For electroporations that included co-expression of dominant negative p53 peptide, GSE56 (CellScript) was added immediately prior to electroporation at a final concentration of 500 nM. Electroporated cells were recovered in complete medium. Genomic DNA (gDNA) was extracted on the indicated days using a gDNA extraction kit (New England Biolabs, T3010) or crude lysis buffer (10 mM Tris pH 8.0, 2 mM EDTA, 0.2% Triton X-100, and 200 μg/mL proteinase K) then analyzed by next-generation sequencing for editing efficiency using CRIS.py^62^. For crude gDNA extraction, samples were incubated at 65 °C for 15 minutes followed by incubation at 95 °C for 5 minutes and diluted 1:10 with water prior to PCR. Sequences of sgRNAs and a list of primers are provided in **Supplementary Tables 1 and 2**, respectively.

### Flow cytometry

Antibodies and dilutions used for flow cytometry are provided in **Supplementary Table 3**. For monitoring apoptosis, DAPI and Apotracker Green was diluted (1:200) and cells were stained per the manufacturer s recommendation (BioLegend, 427403). To perform cell cycle analysis, a cell lysis solution (0.1% w/v sodium citrate, 0.1% Triton X-100, 2 mg/mL RNase A) containing propidium iodide (PI; 0.05 mg/mL) was used. To quantify mitotic cells, phospho-histone H3 (Ser10) was diluted (1:200) in the PI-containing lysis solution. Representative gating strategies are show in **Extended Data Figure 6a, b**. The MicroFlow In Vitro250/50 Kit (Litron Laboratories) was used for micronuclei quantification according to the manufacturer s instructions. After cell harvest, apoptotic and necrotic cells were labeled with the photoactivated dye ethidium monoazide and then washed. After lysis, healthy nuclei and micronuclei were then stained with SYTOX Green. Micronuclei were distinguished from nuclei based on 1/10^th^ – 1/100^th^ DNA-associated fluorescence intensity. Percent micronuclei were calculated by dividing the number of events in the micronuclei gate by the events within the nuclei gate and multiplying by 100. Sample acquisition was performed using a BD FACSymphony A3 or FACSCelesta (BD Biosciences). Data were collected using BD FACS Diva (v.9) and analyzed with FlowJo (v.10).

### Protein extraction and Western blot

Antibodies and dilutions used for Western blot analysis are provided in **Supplementary Table 4**. Cells were washed with DPBS then lysed in Laemmli sample buffer containing β-mercaptoethanol (Bio-Rad, 1610737). Cell lysates were fractionated by electrophoresis on a 4%–15% polyacrylamide gel (Bio-Rad, 4568086). Proteins were transferred to a polyvinylidene fluoride membrane and membranes were incubated in blocking buffer (Bio-Rad, 12010020) for 1 hour at room temperature, then incubated with primary antibody at 4 °C overnight. The membranes were then washed in Tris-buffered saline with 0.1% Tween-20 buffer, incubated with secondary antibody for 1 hour at room temperature, washed again and analyzed using a Bio-Rad ChemiDoc Go imaging system. Data were collected and processed using Image Lab software (v.6).

### Lentiviral vector production

Lentiviral vectors were produced in HEK293 cells grown in high glucose DMEM (Thermo: 11965084) supplemented with 10% FBS, 1x GlutaMAX (Thermo: 35050061), and 100 μM sodium pyruvate (Thermo, 11360070). One day before transfection with vector plasmids, 3.8x10^6^ HEK293T cells were plated into 10 cm plates overnight. Thirty minutes before transfection, 1.64 pmol of pLenti CMV H2B-mNeonGreen (gift from D. Pellman) was mixed with 1.3 pmol of psPAX2 (Addgene Plasmid #12260, gag/pol/rev packaging plasmid) and 0.72 pmol of pMD2.G (Addgene Plasmid #12259, VSV-G envelope plasmid) in 1.5 mL of OPTI-MEM (Thermo, 31985070) and added to 1.5 mL of OPTI-MEM mixed with 50 µl of Lipofectamine 2000 Transfection Reagent (Thermo Fisher Scientific, 11668027). Mixtures were added to HEK293 cells, which were then incubated at 37 °C in 5% CO2 for 18 hours. Fresh medium was then placed on the cells, and viral vector supernatants were collected 48 hours after transfection. Viral supernatant was centrifuged at 500 x *g* for 5 minutes and filtered through a 0.45 μm PES filter. The filtered viral supernatant was concentrated 100x in X-VIVO® 10 medium after centrifugation at 100,000 x *g* for 90 minutes at 4 °C. Serial dilutions of lentiviral vectors were used for transduction of HEK293 cells to calculate titer.

### Live-cell imaging

HSPCs were thawed into complete pre-stimulation medium described above supplemented with GlutaMAX, SR1, and UM729. Twenty-four hours after thawing, cells were transduced with a lentiviral H2B-GFP reporter at a multiplicity of infection of 100. Transduction was performed at 2x10^6^ cells/mL with 10 μM PGE2 (Cayman Chemicals, 14010) and 1 mg/mL LentiBOOST (Sirion Biotech, SB-A-LF-900-01)^63^. Twenty-four hours later, cells were electroporated with CRISPR-Cas9 ribonucleoproteins targeting the *BCL11A* +58 enhancer. On the day of electroporation, ibiTreat μ-Slide 8 well chambers (ibidi, 80806) were coated overnight at 4 °C with 5 μg/mL biotinylated anti-human CD43 antibody (Thermo, MA1-19495, RRID:AB_1072364) diluted in DPBS^64^. Twenty-four hours after genome editing, GFP-positive HSPCs were purified by fluorescence-activated cell sorting (FACSAria, BD Biosciences), resuspended at 5x10⁵ cells/mL, and seeded into antibody-coated imaging chambers. Cells were allowed to adhere and recover overnight before imaging. Time-lapse imaging was performed for 72 hours using a Marianas live-cell imaging system composed of a Yokogawa CSU-X Spinning Disc and a Photometric Prime 95B camera (3i, Intelligent Imaging Innovations) equipped with a Zeiss 100x/1.4 NA oil immersion objective in a humidified environmental chamber maintained at 37 °C and 5% CO₂. Images were acquired every 20 minutes using SlideBook 6 software (Intelligent Imaging Innovations) with Z-stack acquisition (10 μm stack with 500 nm interval) and Definite Focus 2 autofocus. GFP-positive HSPCs containing micronuclei were identified by manual inspection of each well and tracked throughout the imaging period to assess mitotic progression and cell fate. Representative still images and time-lapse videos were generated from Z-stack image sequences using Imaris (Oxford Instruments).

### Immunofluorescence microscopy and image analysis

Antibodies and dilutions used for immunofluorescence are provided in **Supplementary Table 4**. To bypass G2/M checkpoint arrest and enrich mitotic cells for visualization of BUB1 localization at centromeres, HSPCs were treated with the ATR inhibitor VE-821 (MedChemExpress, HY-14731) at 10 μM for 2.5 hours in culture medium. Cells were then collected, resuspended in pre-warmed PHEM buffer (EMS, 11163) containing 10 μM VE-821, and allowed to settle onto glass coverslips for 30 minutes at room temperature, resulting in a total VE-821 exposure of 3 hours. Cells were fixed with 4% methanol-free paraformaldehyde (EMS, 15700) in PHEM buffer for 20 minutes at room temperature, washed with PHEM, and permeabilized with 0.5% Triton X-100 in PHEM for 20 minutes. Cells were then blocked for 1 hour at room temperature in 5% normal donkey serum (Jackson Immuno Research Labs, 017000121) prepared in PHEM containing 0.1 M glycine and 0.1% Triton X-100. Primary antibodies were diluted in blocking buffer and incubated for 1 hour at room temperature followed by overnight incubation at 4 °C. After being washed with 0.1% Triton X-100 in PHEM buffer, coverslips were incubated with species-appropriate fluorophore-conjugated secondary antibodies for 2 hours at room temperature in the dark. Rabbit anti-human H2A-T120p antibody was conjugated to CoraLite Plus 647 (Proteintech, KFA503) according to the manufacturer s protocol. After secondary antibody staining and washing, coverslips were incubated with the conjugated anti-rabbit H2A-T120p antibody for an additional 1 hour. Nuclei were counterstained with Hoechst 33342 (10 μg/mL, Thermo, H3570) diluted in PHEM for 10 minutes, and coverslips were mounted using ProLong Glass Antifade mountant (Thermo Fisher Scientific) and cured for 72 hours before imaging.

Images were acquired using a Nikon Ti2 inverted confocal field scanning spinning disk microscope equipped with a plan apochromat 100x/1.45 NA oil immersion objective and the ORCA Fusion BT sCMOS camera. The system has a 200-micron range Piezo stage that was used for high-precision z-axis positioning and speed. Excitation was achieved using 405, 488, 561, and 640 nm laser lines. Microscope control and image acquisition were performed using NIS-Elements Advanced Research software (v.6). Acquisition parameters, including laser power, exposure time, camera gain, and pixel size, were maintained across all samples within the experiment with 40% laser power and 300 ms exposure time for the 405 nm channel, 50% laser power and 400 ms exposure time for the 488 nm channel, 60% laser power and 600 ms for the 561 nm channel, and 70% laser power and 700 ms exposure time for the 640 nm channel. Z-stacks encompassing the full cellular volume were collected with a step size of 0.2 μm.

Raw image datasets were processed in NIS-Elements Advanced Research software (v.6). Three-dimensional visualization of acquired Z-stacks was performed using alpha-blended volume rendering. For quantitative analysis of BUB1 and H2A-T120 fluorescence associated with CREST-defined centromeric regions, all nuclei were first identified using a binary mask generated from the Hoechst channel. Mitotic cells were identified using an object-based segmentation pipeline that incorporated sphericity and volume. Individual object masks were subsequently generated for BUB1, CREST, and H2A-T120p signals detected in the 488, 561, and 640 nm channels, respectively. Colocalized BUB1, CREST, and H2A-T120p puncta located within mitotic cells were identified, and the mean fluorescence intensity of BUB1 (488) and H2A-T120p (647) within these overlapping objects was measured and recorded for comparison across samples.

### Alkaline Comet Assay

DNA strand breaks were quantified using the CometAssay HT Kit and CometAssay Electrophoresis System II (R&D Systems/Bio-Techne) according to the manufacturer’s protocol with minor modifications. Briefly, 10,000 cells were resuspended in 20 μL PBS, and 10 μL of the suspension was mixed with 90 μL molten low-melting-point agarose maintained at 37 °C. Approximately 50 μL of the cell-agarose mixture (2,500 cells) was applied to each well of a CometSlide and allowed to solidify for 10 minutes at room temperature. Slides were then immersed in pre-chilled lysis solution and incubated overnight at 4 °C. Slides were transferred to freshly prepared alkaline unwinding buffer (200 mM NaOH, 1 mM EDTA; pH >13) for 1 hour at 4 °C before electrophoresis in alkaline electrophoresis buffer (200 mM NaOH, 1 mM EDTA; pH >13) at 21 V for 30 minutes using the CometAssay Electrophoresis System II maintained at 4°C. After electrophoresis, slides were washed twice in distilled water for 5 minutes each, fixed in 70% ethanol for 5 minutes, and air dried. DNA was stained with 1X SYBR Gold (Thermo, S11494) for 30 minutes at room temperature in the dark, rinsed with distilled water, and dried before imaging. Images were acquired using an EVOS fluorescence microscope (Thermo Fisher Scientific). Representative low-magnification images were first collected to identify the agarose boundaries, after which approximately five non-overlapping fields near the center of each agarose well were imaged using a 10X objective with identical acquisition settings across all samples within an experiment. Comets were quantified using the OpenComet v2 plugin for ImageJ/Fiji^65^. Automated comet identification was followed by manual inspection of segmentation overlays to exclude debris, overlapping nuclei, and incorrectly segmented comets before final data export. DNA damage was quantified using the tail moment calculated by OpenComet.

### Xenotransplantation of gene-edited CD34^+^ HSPCs into NBSGW mice

Female NBSGW mice aged 6–8 weeks were purchased from The Jackson Laboratory (stock #: 026622, RRID:IMSR_JAX:026622). Busulfan was administered (10 mg/kg) as an intraperitoneal injection 48 h before injection of human HSPCs. HSPCs were treated with Q-VD-OPh (1 μM) immediately after electroporation for 24 hours. Unedited, edited, and Q-VD-OPh–treated edited HSPCs were collected, washed with DPBS, and resuspended in DPBS+2% FBS for retro-orbital injection. HSPCs were administered at a dose of 2.5x10^5^ cells per mouse. Chimerism and editing efficiency were evaluated at 8 weeks post-transplantation in the bone marrow at the time of euthanasia. Cell lineage composition was determined in the bone marrow using mouse- and human-specific antibodies (**Supplementary Table 3**). Representative gating strategies are show in **Extended Data Figure 6c**. Sample acquisition was performed using a Cytek Aurora. Data were collected using SpectroFlo (v.4) and analyzed with FlowJo (v.10).

### Statistical analysis

All statistical analyses were performed using GraphPad Prism (v.10). Data are presented as mean ± SD, with individual biological replicates shown where possible. Biological replicates represent independent HSPC donors, and technical replicates were averaged prior to statistical testing. HSPCs from the same donor were distributed across all experimental conditions; therefore, paired statistical tests were used to account for the donor-matched experimental design and inter-donor biological variability. For comparisons between two groups, paired two-tailed Student s t-tests were used except for the xenotransplant data for which an unpaired t-test was used. For comparisons involving more than two groups at a single time point, one-way analysis of variance (ANOVA) with Holm-Šídák’s multiple comparisons test was applied. For experiments involving multiple groups measured over time or across conditions, repeated-measures two-way ANOVA with the Geisser-Greenhouse correction was used, with donor treated as a matched factor for repeated-measures analyses. Main effects (e.g., time and treatment) and interaction terms are reported for all models. Post hoc multiple comparisons were performed using Tukey’s multiple comparisons test. Statistical significance was defined as p < 0.05. No statistical methods were used to predetermine sample size. Sample sizes were chosen based on previous experience with similar experimental systems. Except for confocal immunofluorescence imaging, investigators were not blinded to group allocation during experiments or outcome assessment.

## Supporting information

Unprocessed Western blots

## DATA AVAILABILITY

The source data underlying all graphs in this study are provided in a Source Data file. Raw data files are available from the lead contact upon reasonable request.

## ACKNOWLEDGMENTS

The authors would like to thank M. Leibowitz, D. Pellman, R. Sharma, E. Potchanant, and L. Kabeche for valuable feedback and technical advice. The authors would like to thank Versiti Blood Research Institute Shared Resources (RRID, SCR_025503) and Wisconsin-Madison University Biotechnology Center DNA Sequencing Core Facility (RRID:SCR_017759) for their services, access to instrumentation, and specialist support. The authors thank the members of the St. Jude Children’s Research Hospital Light Microscopy, Hartwell Center, Center for Advanced Genome Engineering, and Flow Cytometry and Cell Sorting Shared Resources. This research was supported by the National Institute of Diabetes and Digestive and Kidney (NIDDK) Diseases of the National Institutes of Health (NIH) from a Cooperative Centers of Excellence in Hematology (CCEH) Pilot & Feasibility award U24 DK126127 (PAD) and through use of the Hematopoietic Cell Procurement and Processing Resource at Fred Hutchinson Cancer Center. This work was supported by NIH grants K01 DK132453 (PAD), R03 DK142988 (PAD), and the Versiti Blood Research Institute Foundation (VBRIF; PAD). The CCEH Cell Core is supported by NIH grant U54 DK106829, and the St. Jude Shared Resource Laboratories are supported in part by ALSAC and the NIH National Cancer Institute grant P30 CA21765. The content is solely the responsibility of the authors and does not necessarily represent the official views of the funders. The funders had no role in the study design, the data collection and analysis, the decision to publish the work, or the preparation of the manuscript.

## AUTHOR CONTRIBUTIONS

Z.B., S.S., M.H., and K.D.M. performed experiments. J.G.M. provided expertise in immunofluorescence analysis. R.B.D. provided expertise in micronucleus flow cytometry. A.P. provided expertise in live-cell imaging. P.A.D. designed the study, performed and interpreted experiments, analyzed the data, assembled the figures, and wrote the manuscript. All authors reviewed and approved the final manuscript.

## EXTENDED DATA FIGURES AND LEGENDS

**Extended Data Figure 1.**
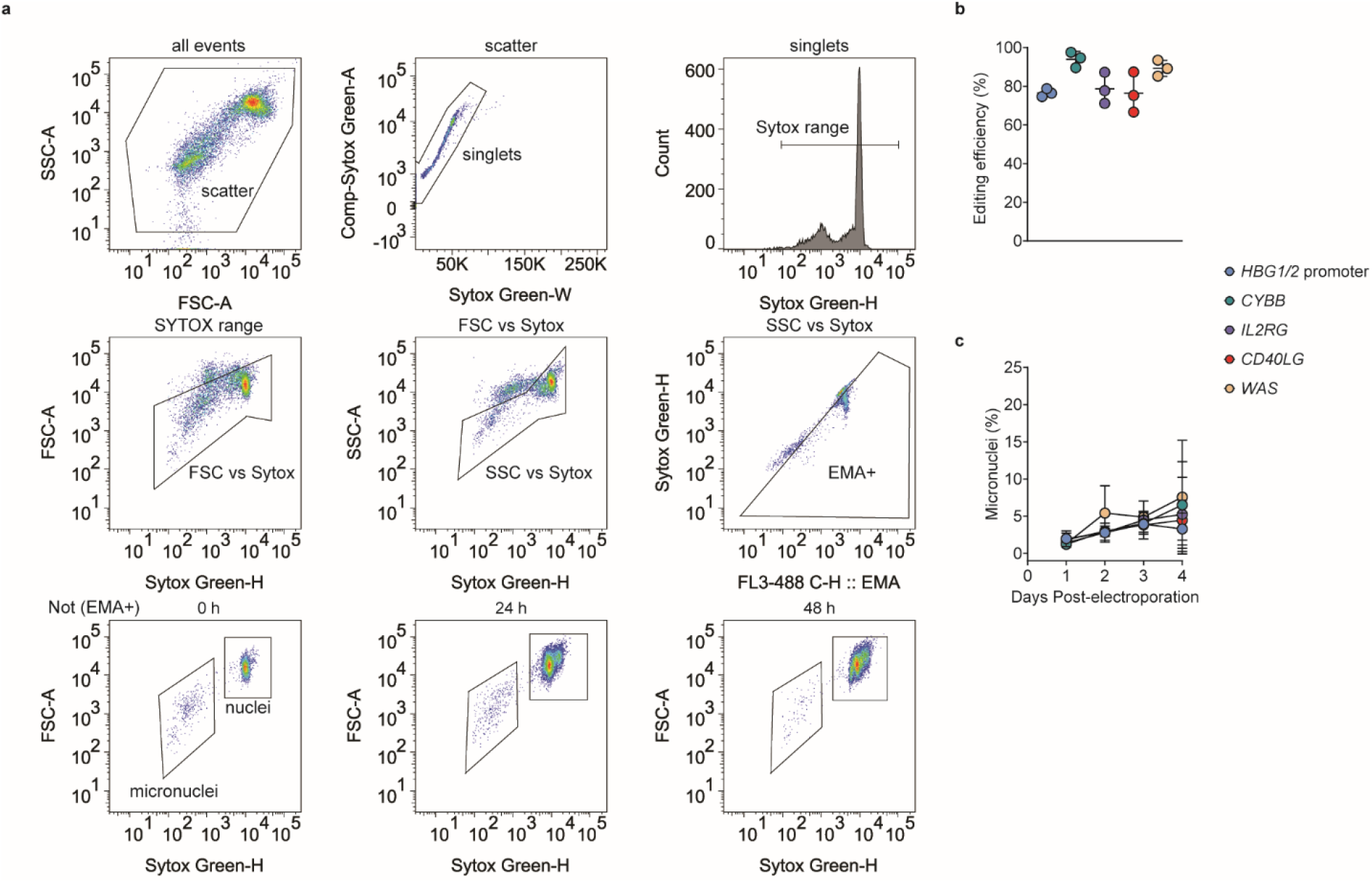
Micronucleus formation occurs across multiple genome editing targets in HSPCs. **a**, Gating strategy used to identify micronuclei. Ethidium monoazide (EMA) labels dead or dying cells for exclusion and SYTOX green labels viable nuclei and micronuclei after lysis. The micronuclei gate was defined based on fluorescence intensity approximately 1/10 to 1/100 that of the nuclei population, according to the manufacturer’s protocol. **b,** Genome editing efficiency determined by next-generation sequencing at therapeutic loci on day 7 post-electroporation. **c,** Flow cytometry for micronuclei on days 1–4 post-electroporation. For b and c, HSPCs were pre-stimulated for 48 h before electroporation of Cas9 RNP targeting the relative locus using a MaxCyte GTx. Mean ± standard deviation. *n* = 3 donors performed in triplicate.

**Extended Data Figure 2.**
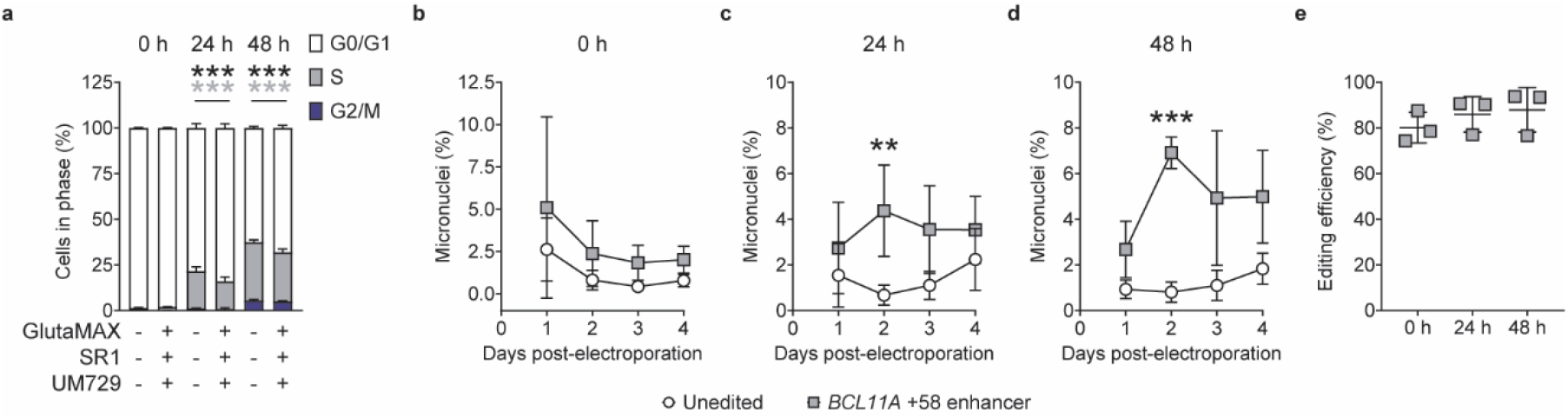
Temporal changes in cell-cycle distribution, micronucleus frequency, and editing efficiency. **a,** Cell-cycle distribution of HSPCs without pre-stimulation (0 h) or with pre-stimulation for 24 or 48 h ± GlutaMax/SR1/UM729. Mean ± standard deviation with a two-way ANOVA. *n* = 3 donors. Tukey’s multiple comparisons test \*\*\**P* < 0.001. Black *** indicates significant differences in G0/G1 phase and grey *** indicates significant differences in S phase. **b–d,** Flow cytometry for micronuclei on days 1–4 post-electroporation in HSPCs without pre-stimulation (0 h) or with pre-stimulation for 24 or 48 h in cultures without GlutaMax/SR1/UM729. Mean ± standard deviation with a two-way repeated measures ANOVA with the Geisser-Greenhouse correction. *n* = 3 donors performed in triplicate. Treatment effect: 0 h, *P* = 0.2908; 24 h, *P* = 0.1006; 48 h, *P* = 0.0029. Tukey’s multiple comparison test \*\**P* < 0.01, \*\*\**P* < 0.001. **e**, Genome editing efficiency of cultures in b–d determined by next-generation sequencing on day 7 post-electroporation. Mean ± standard deviation. *n* = 3 donors performed in triplicate.

**Extended Data Figure 3.**
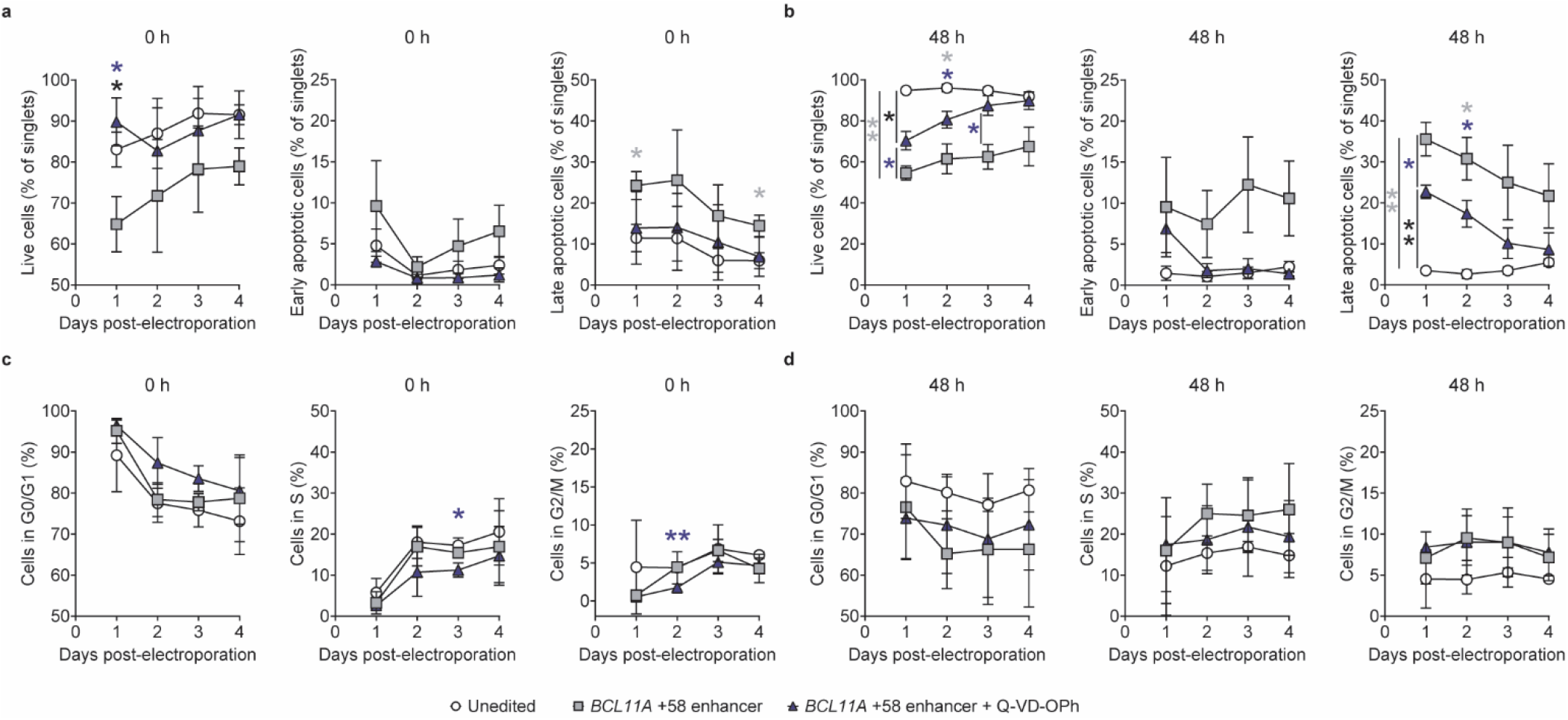
Caspase inhibition alters apoptosis and cell cycle distribution after genome editing in HSPCs. **a,** Flow cytometry analysis of apoptosis on days 1–4 post-electroporation in HSPCs without pre-stimulation (0 h) before electroporation of Cas9 RNP targeting the *BCL11A* +58 enhancer ± Q-VD-OPh. Panels show the frequency of live (left), early apoptotic (middle), and late apoptotic (right) cells. Mean ± standard deviation with a two-way repeated measures ANOVA with the Geisser-Greenhouse correction. *n* = 3 donors performed in triplicate. Treatment effect: live cells, *P* = 0.0499. Tukey’s multiple comparisons test \**P* < 0.05. Black * on live cells measured on day 1 indicates significant differences between unedited and edited. Blue * on live cells measured on day 1 indicates significant differences between edited and edited + Q-VD-OPh. Grey * on late apoptotic cells measured on days 1 and 4 indicate significant differences between unedited and edited. **b,** Flow cytometry analysis of apoptosis on days 1–4 post-electroporation in HSPCs with 48 h pre-stimulation before electroporation of Cas9 RNP targeting the *BCL11A* +58 enhancer ± Q-VD-OPh. Panels show the frequency of live (left), early apoptotic (middle), and late apoptotic (right) cells. Mean ± standard deviation with a two-way repeated measures ANOVA with the Geisser-Greenhouse correction. *n* = 3 donors performed in triplicate. Treatment effect: live cells, *P* = 0.0003. Tukey’s multiple comparisons test \**P* < 0.05, \*\**P* < 0.01. Significant differences between groups on live cells and late apoptotic cells measured on day 1 are indicated by lines. Grey * and blue * on live cells measured on day 2 indicate significant differences between unedited vs. edited and unedited vs. edited + Q-VD-OPh, respectively. Grey * and blue * on live cells measured on day 3 indicate significant differences between unedited vs. edited and edited vs. edited + Q-VD-OPh, respectively. Grey * and blue * on late apoptotic cells measured on day 2 indicate significant differences between unedited vs. edited and unedited vs. edited + Q-VD-OPh, respectively. **c,** Flow cytometry analysis of cell cycle distribution on days 1–4 post-electroporation in HSPCs without pre-stimulation (0 h) before electroporation of Cas9 RNP targeting the *BCL11A* +58 enhancer ± Q-VD-OPh. On the left is G0/G1 phase, in the middle is S phase, and on the right is G2/M phase. Mean ± standard deviation with a two-way repeated measures ANOVA with the Geisser-Greenhouse correction. *n* = 3 donors performed in triplicate. Tukey’s multiple comparisons test \**P* < 0.05, \*\**P* < 0.01. Blue * on S phase measured on day 3 indicates significant differences between unedited and edited + Q-VD-OPh. Blue ** on G2/M phased measured on day 2 indicates significant differences between edited and edited + Q-VD-OPh. **d,** Flow cytometry analysis of cell cycle distribution in HSPCs pre-stimulated for 48 h before electroporation of Cas9 RNP targeting the *BCL11A* +58 enhancer ± Q-VD-OPh. On the left is G0/G1 phase, in the middle is S phase, and on the right is G2/M phase. Mean ± standard deviation. *n* = 3 donors performed in triplicate.

**Extended Data Figure 4.**
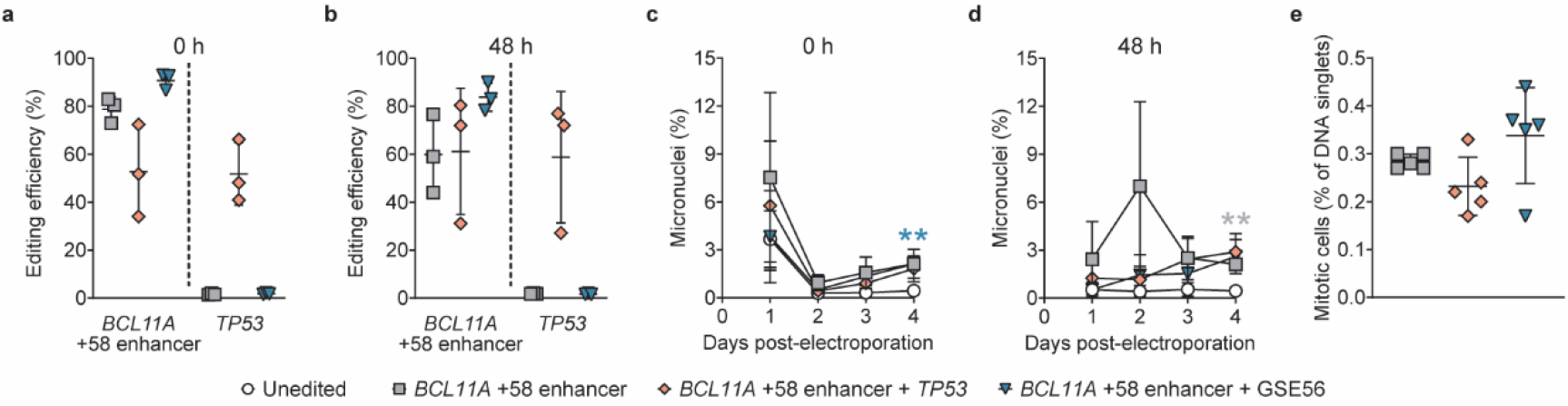
Effects of p53 suppression on micronucleus formation and mitotic cells after genome editing. **a,** Genome editing efficiency in HSPCs without pre-stimulation (0 h) before electroporation of Cas9 RNP targeting the *BCL11A* +58 enhancer, combined editing at the *BCL11A* +58 enhancer and *TP53*, or *BCL11A* +58 enhancer with GSE56 determined by next-generation sequencing (NGS) on day 7 post-electroporation. Mean ± standard deviation. *n* = 3 donors performed in triplicate. **b,** Genome editing efficiency in HSPCs with 48 h pre-stimulation before electroporation of Cas9 RNP targeting the *BCL11A* +58 enhancer, combined editing at the *BCL11A* +58 enhancer and *TP53*, or *BCL11A* +58 enhancer with GSE56 determined by NGS on day 7 post-electroporation. Mean ± standard deviation. *n* = 3 donors performed in triplicate. **c,** Flow cytometry for micronuclei on days 1–4 post-electroporation of cultures in a. Mean ± standard deviation with a two-way repeated measures ANOVA with the Geisser-Greenhouse correction. *n* = 3 donors performed in triplicate. Time effect: *P* = 0.0049. Treatment effect: *P* = 0.1988. Tukey’s multiple comparisons test \*\**P* < 0.01. Teal ** indicates significant differences between unedited and edited + GSE56. **d,** Flow cytometry for micronuclei on days 1–4 post-electroporation of cultures in b. Mean ± standard deviation with a two-way repeated measures ANOVA with the Geisser-Greenhouse correction. *n* = 3 donors performed in triplicate. Time effect: *P* = 0.3647. Treatment effect: *P* = 0.0001. Tukey’s multiple comparisons test \*\**P* < 0.01. Grey ** indicates significant differences between unedited and edited. **e,** Flow cytometry analysis for phospho-histone H3-S10-postive cells 1-day post-electroporation in HSPCs with 48 h pre-stimulation. Mean ± standard deviation. *n* = 5 donors.

**Extended Data Figure 5.**
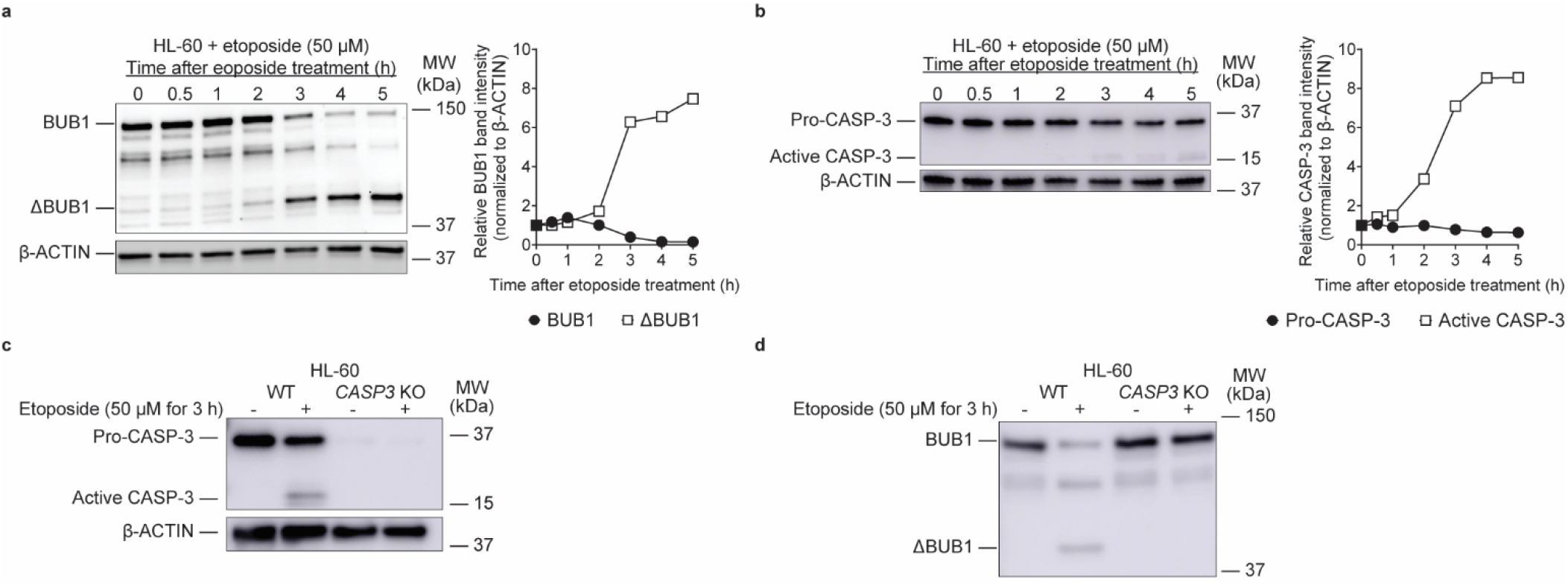
CASPASE-3-dependent BUB1 cleavage following etoposide treatment in HL-60 cells. **a,** Immunoblot for BUB1 in HL-60 cells treated with etoposide (left). HL-60 cells were treated with 50 μM etoposide and collected at the indicated times. Quantification of BUB1 fragments normalized to β-ACTIN (right). **b,** Immunoblot for CASPASE-3 in cultures as in a (left). Quantification of pro- and cleaved CASPASE-3 normalized to β-ACTIN (right). **c,** Immunoblot for CASPASE-3 and β-ACTIN in wild-type (WT) and bulk *CASP3* edited HL-60 cells treated with 50 μM etoposide. **d,** Immunoblot for BUB1 and β-ACTIN in wild-type (WT) and bulk *CASP3* edited HL-60 cells treated with 50 μM etoposide.

**Extended Data Figure 6.**
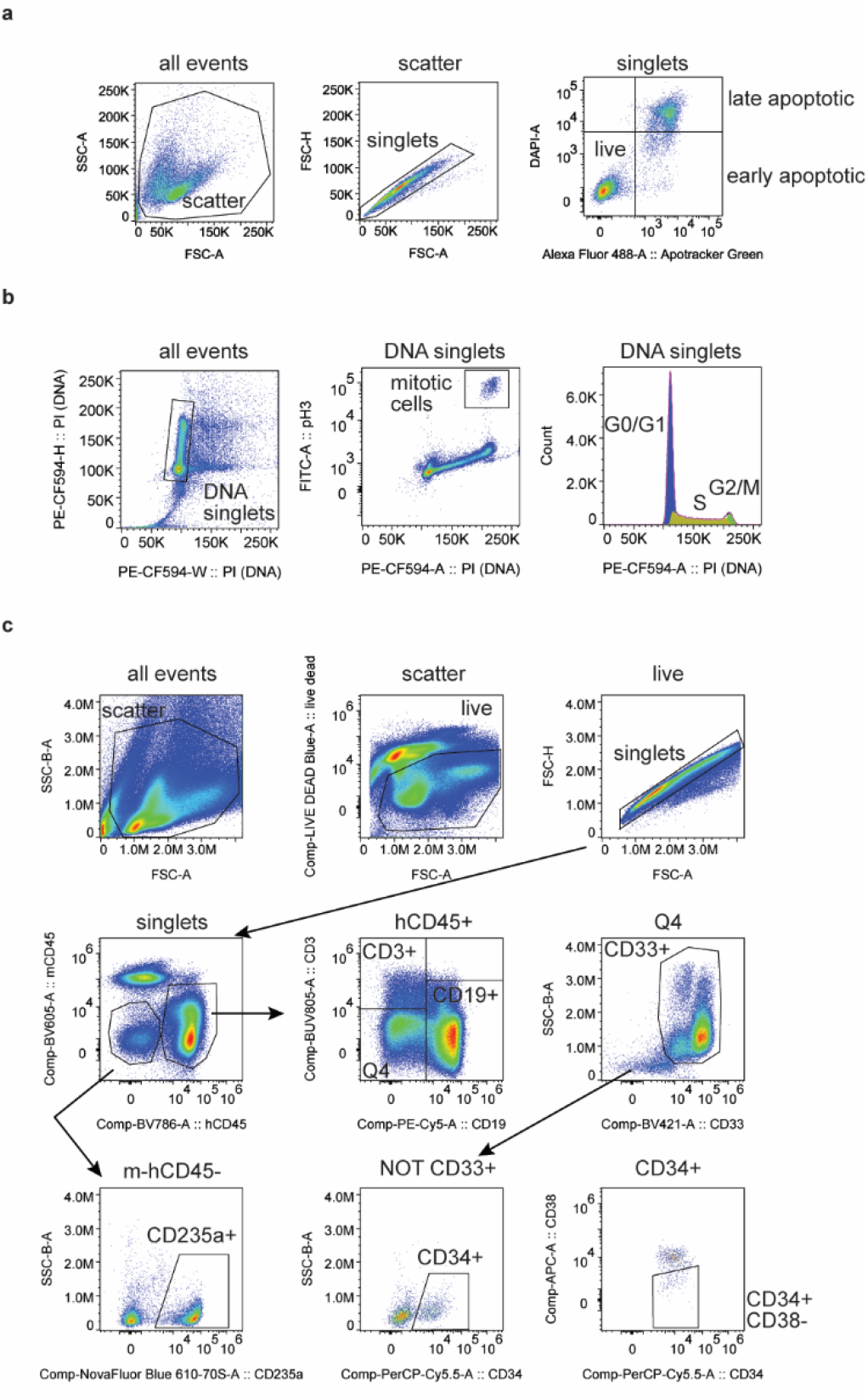
Flow cytometry gating strategies. **a,** Gating strategy for calculating live cells, early apoptotic cells, and late apoptotic cells. **b,** Gating strategy for calculating mitotic cells and cell cycle distribution analysis. **c,** Gating strategy for calculating mouse and human bone marrow cell populations. All antibodies and reagents are described in the methods section.

**Supplemental Movies 1–3.** Live cell imaging videos of genome-edited CD34^+^ HSPCs with micronuclei as in Fig. 1f, g (H2B-mNeonGreen). Timestamp shows relative time in h:m:s.ms. Confocal imaging under a 100x objective. Dynamic scale bar adapts to digital zoom. Single video frames have been extracted and are presented in Fig. 1g.

