## Supplementary material for "Caspase Activity Affects Cell Cycle and Chromosome Integrity after Genome Editing in Human Hematopoietic Stem and Progenitor Cells": Unprocessed Western blots

Source Data

Figure 3a

α-TUBULIN (top band) and CASPASE-3 (bottom band) - all 4 donors

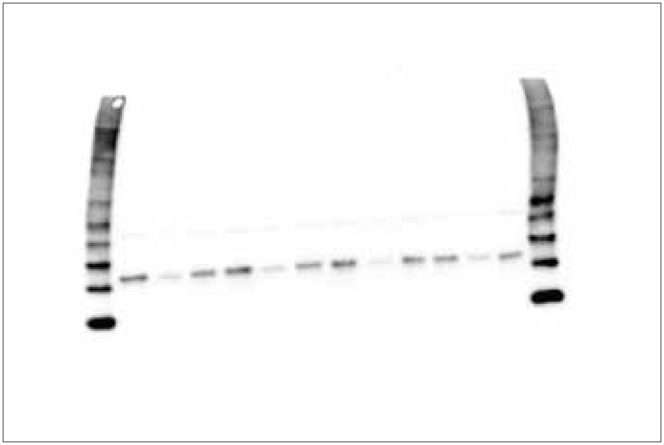

Figure 3c

BUB1 (top bands) and β-ACTIN (bottom band) - all 4 donors

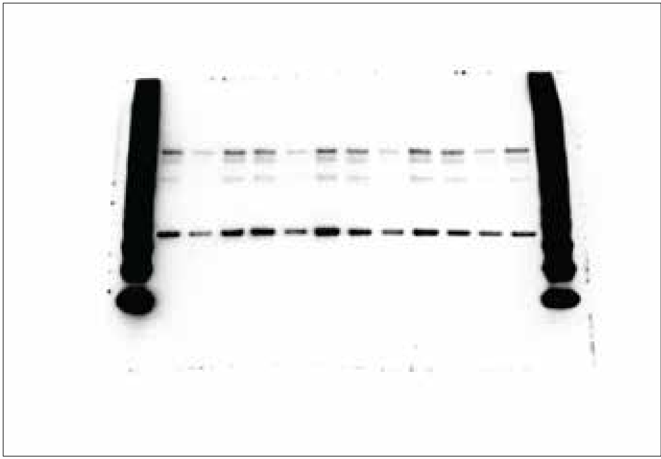

Figure 3l

BUB1 (top band) and β-ACTIN (bottom band) - donors 1 and 2

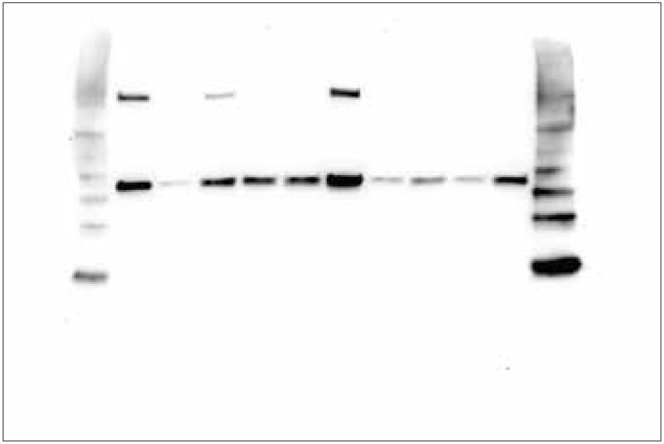

BUB1 (top band) and β-ACTIN (bottom band) - donors 3 and 4

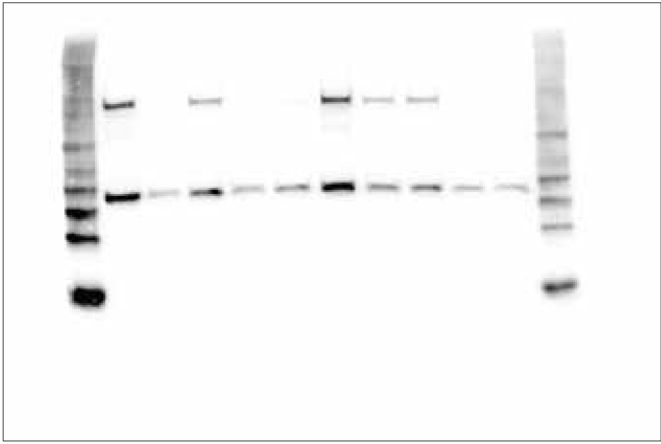

Extended Data Figure 5a

BUB1

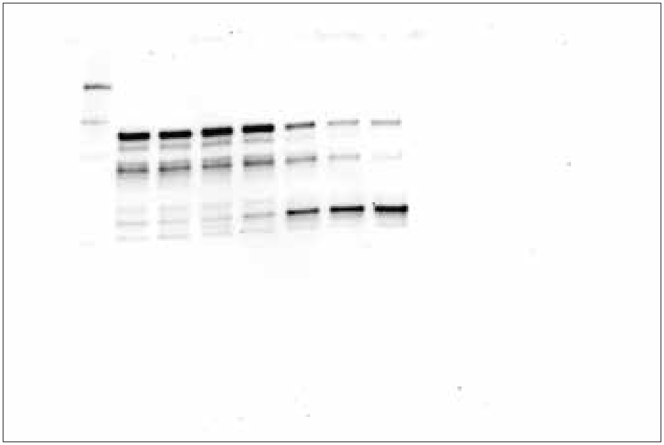

β-ACTIN

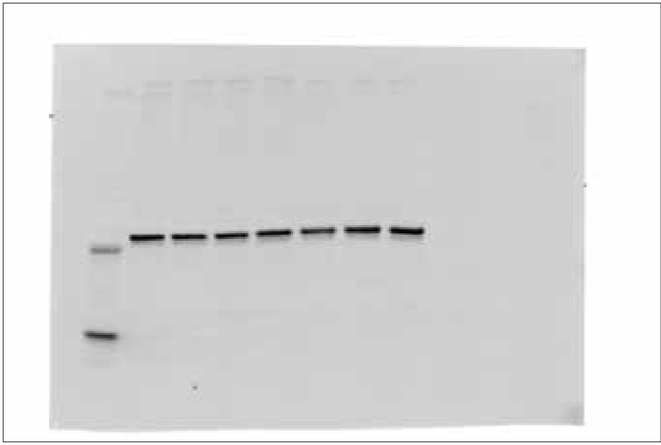

Extended Data Figure 5b

CASPASE-3

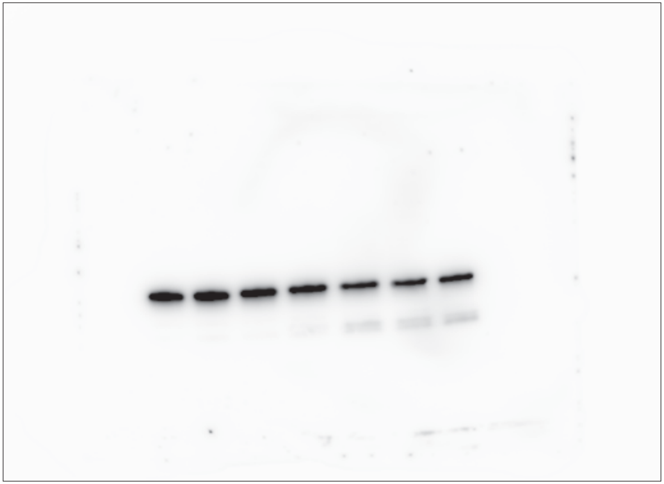

β-ACTIN

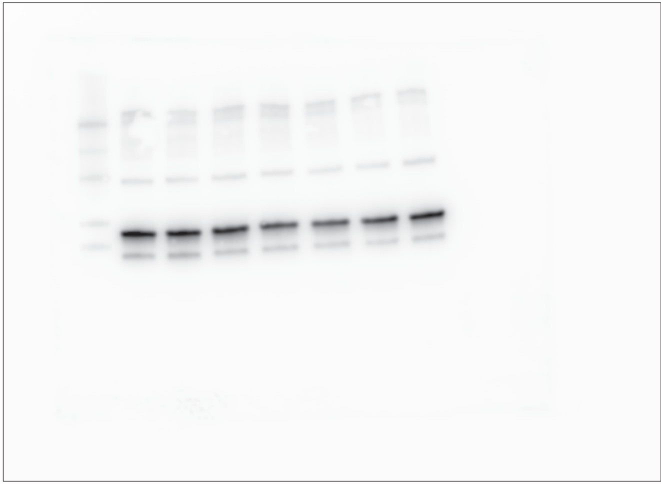

**Extended Data Figure 5c**

CASPASE-3

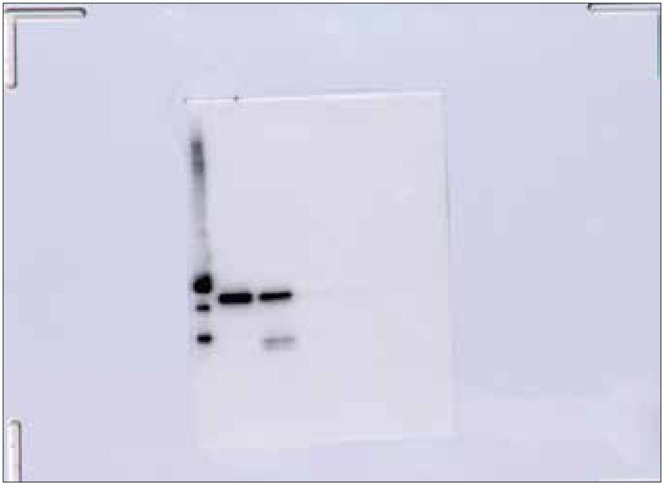

β-ACTIN

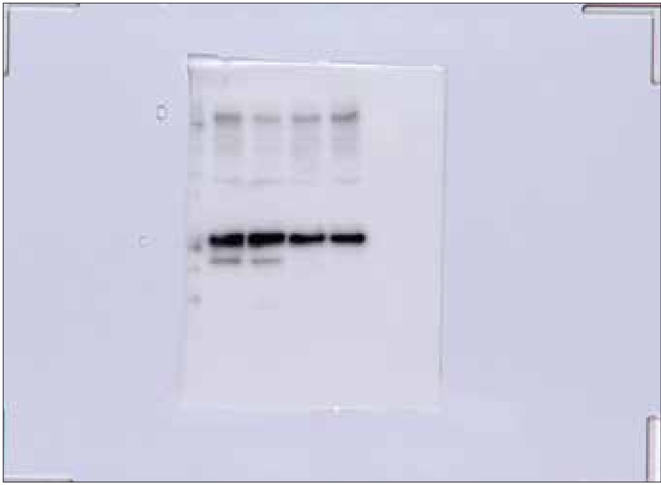

**Extended Data Figure 5d**

BUB1

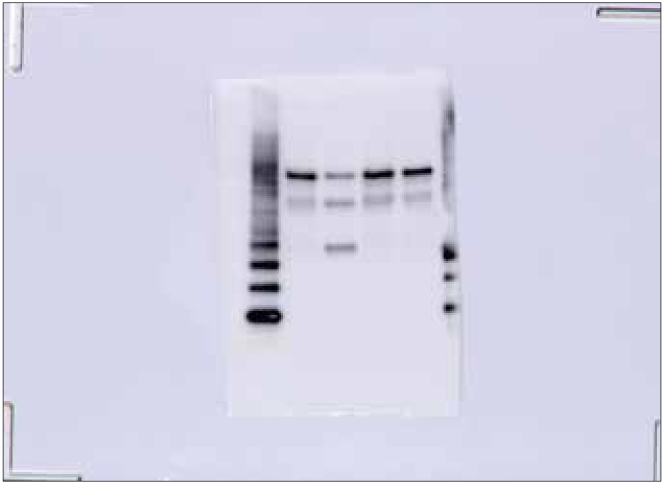
