## Supplemental tables for "Caspase Activity Affects Cell Cycle and Chromosome Integrity after Genome Editing in Human Hematopoietic Stem and Progenitor Cells"

| **Name** | **Sequence (without PAM)** |
|---|---|
| *BCL11A* + 58 Enh | CTAACAGTTGCTTTTATCAC |
| *HBG1/2* Pro | CTTGTCAAGGCTATTGGTCA |
| *IL2RG* | TGGTAATGATGGCTTCAACA |
| *CYBB* | TTGTGCAGCTGGTTTGGCTG |
| *CD40LG* | GTATCTTCTGGCAGAGAAGG |
| *WAS* | AGGGCAGAAAGCACCATGAG |
| *TP53* | TCCTCAGCATCTTATCCGAG |
| *CASP3* | TGTCGATGCAGCAAACCTCA |
| *BUB1* | GCCAGTGCTGTCCTTCAGAG |

| **Name** | **Forward Sequence** | **Reverse Sequence** |
|---|---|---|
| *BCL11A* + 58 Enh | GATACAGGGCTGGCTCTATGC | CAAGAGAGCCTTCCGAAAGAGG |
| *HBG1/2* Pro | ACTGAATCGGAACAAGGCAAAGGCT | ACCCATGGCGTCTGGACTAGGAGCT |
| *IL2RG* | AGGACTTAGCCCGTGTCACACAGCA | CCCCTTCCCACAGCCACCCTTCTCA |
| *CYBB* | ACACTGTTGAGGAAAGAGCTGGCTC | GCAATGGAGGGAGTGAGGCTAATGGT |
| *CD40LG* | GCAACGATTGTGCGCTCTTA | GGCTTGTGGTTCATCTTACCT |
| *WAS* | GGGCCCAAGCTCAGCCTAACGAGGA | GGGCAGGAGATCCCCAGCTCACCAA |
| *TP53* | GGCGCTGCCCCCACCATGAG | CTGGAGGGCCACTGACAACCACCCT |
| *CASP3* | TCACTACCTAAACCGCTGTG | TTGTTTCTTCTTACCATCACGC |
| *BUB1* | TCATCACAGAGAGTTTGACTTTGT | TTGGGTTCGTGTAGGCTGAG |

**Forward-prefix**: CACTCTTTCCCTACACGACGCTCTTCCGATCT (32bp)

**Reverse-prefix**: GTGACTGGAGTTCAGACGTGTGCTCTTCCGATCT (34bp)

| Target | Fluorochrome | Clone | Concentration (ug/mL) | Dilution | Final Concentration (ug/mL) | Manufacturer | Cat# | RRID | Isotype | Concentration ug/mL | Dilution to Match Target Final Concentration | Manufacturer | Cat# | RRID |
|---|---|---|---|---|---|---|---|---|---|---|---|---|---|---|
| phospho-H3S10 | Alexa Fluor 488 | D2C8 | | 200 | | Cell Signaling Technologies | 3465 | AB_10695860 | IgG | | | | | |
| live/dead blue | | | | 100 | | Thermo | L23105 | AB_3717566 | | | | | | |
| CD3 | BUV805 | SK7 | 200 | 50 | 4.000 | BD | 612893 | AB_2870181 | IgG1 | | | | | |
| CD33 | BV421 | P67.6 | 200 | 50 | 4.000 | BD | 744761 | AB_2742462 | IgG1 | 200 | 50 | BD | 562438 | AB_11207319 |
| mCD45 | BV605 | 30-F11 | 100 | 100 | 1.000 | BioLegend | 103140 | AB_2562342 | IgG2b | 100 | 100 | BioLegend | 400650 | AB_2864282 |
| hCD45 | BV786 | HI30 | 100 | 100 | 1.000 | BD | 563716 | AB_2716864 | IgG1 | 200 | 200 | BD | 563330 | AB_2869484 |
| CD235a | NovaFluorBlue610-70S | GA-R2 (HIR2) | 100 | 400 | 0.250 | Thermo | H050T03B06 | AB_2925987 | IgG2b | | | | | |
| CD34 | PerCP-Cy5.5 | 581 | 200 | 20 | 10.000 | BioLegend | 343522 | AB_2228973 | IgG1 | 200 | 20 | BioLegend | 400150 | AB_893664 |
| CD19 | PE-Cy5 | HIB19 | 6.3 | 50 | 0.126 | BD | 555414 | AB_395814 | IgG1 | 13 | 103 | BD | 555750 | AB_396092 |
| CD38 | APC | HIT2 | 6.3 | 20 | 0.315 | BD | 555462 | AB_398599 | IgG1 | 13 | 41 | BD | 555751 | AB_398613 |
| | | | | | | | | | | | | iso dilution different from target | | |

| Block | Dilution | Manufacturer | Cat# |
|---|---|---|---|
| CellBlox™ Plus Blocking Buffer | 20 | Thermo | C001T06F01 |
| BD Horizon™ Brilliant Stain Buffer Plus | 5 | BD | 568264 |

| **Target** | **Species** | **Clonality** | **Isotype** | **Supplier** | **Cat.** | **Application** | **Dilution** | **RRID** |
|---|---|---|---|---|---|---|---|---|
| BUB1 | Rabbit | Monoclonal | IgG | abcam | ab195268 | WB, IF | 1000, 100 | AB_3720167 |
| CREST sera | Human | Monoclonal | IgG | Immunovision | HCT-0100 | IF | 1000 | AB_2744669 |
| H2A-p120 | Rabbit | Polyclonal | IgG | Active Motif | 61195 | IF | 1ug/ml | AB_2793548 |
| ACTB | Mouse | Monoclonal | IgG2a | Millipore-Sigma | A2228 | WB | 20000 | AB_476697 |
| CASP3 | Rabbit | Polyclonal | IgG | CST | 9662 | WB | 1000 | AB_331439 |
| Anti-mouse-HRP | Goat | Polyclonal | IgG | Invitrogen | A28177 | WB | 5000 | AB_2536163 |
| Anti-rabbit-HRP | Goat | Polyclonal | IgG | Invitrogen | A27036 | WB | 5000 | AB_2536099 |
| Anti-rabbit-AlexaFluorPlus488 | Donkey | Polyclonal | IgG | Invitrogen | A32790 | IF | 1000 | AB_2762833 |
| Anti-human-DyLight550 | Donkey | Polyclonal | IgG | Invitrogen | SA5-10127 | IF | 2500 | AB_2556707 |
